# Mrc1^Claspin^ is essential for heterochromatin maintenance in *Schizosaccharomyces pombe*

**DOI:** 10.1101/2023.03.28.534615

**Authors:** Kei Kawakami, Yukari Ueno, Nao Hayama, Katsunori Tanaka

## Abstract

Heterochromatin is a highly condensed chromatin structure that represses gene expression. In eukaryotes, maintenance of the heterochromatin structure during cell proliferation is essential for guaranteeing cell identity. However, how heterochromatin is maintained and transmitted to the daughter cells remains largely unknown. In this study, we constructed a reporter system to study the maintenance of heterochromatin in the subtelomeric region of the fission yeast, *Schizosaccharomyces pombe*. We demonstrated that once subtelomeric heterochromatin was established, it tended to be maintained as a metastable structure through cell proliferation. Using this system, we screened an *S. pombe* genome-wide gene deletion library and identified 57 factors required for the maintenance of subtelomeric heterochromatin. We focused on Mrc1^Claspin^, a mediator of DNA replication checkpoint. We found that Mrc1 maintains heterochromatin structure not only at the subtelomeres but also at other heterochromatic loci, such as the pericentromeres and mating-type regions. Furthermore, we showed that Mrc1 facilitates the hypoacetylation state of histone H3K14 by recruiting the Snf2/Hdac-containing Repressor Complex (SHREC), via physical interaction. In addition, depletion of Mst2, an H3K14 acetyltransferase, restored heterochromatin integrity in *mrc1* mutants. This is the first report to show a link between DNA replication factors and H3K14 deacetylation in heterochromatin.

## Introduction

Despite having the same genome, eukaryotes have various types of cells in their bodies, including undifferentiated and differentiated cells. During cell differentiation, eukaryotes alter the expression of individual genes via epigenetic mechanisms, giving rise to cells with new individual characteristics. Programmed gene expression patterns are memorized during proliferation, with cellular memory essential for guaranteeing cellular identity and maintaining tissue and individual homeostasis. Gene expression patterns in eukaryotic cells are regulated by two types of chromatin structure: euchromatin and heterochromatin (1). In euchromatin, histone H3 lysine 9 acetylation (H3K9ac), histone H3 lysine 14 acetylation (H3K14ac), and histone H3 lysine 4 methylation (H3K4me) are epigenetic changes that positively regulate gene transcription. Alternatively, heterochromatin is characterized by histone H3 lysine 9 methylation (H3K9me) and hypoacetylation of H3K14, which negatively regulate gene transcription. The stable maintenance of H3K9me and hypoacetylation of H3K14 is essential for cellular memory (2, 3).

The best-understood mechanism in the process of heterochromatin establishment is RNA interference (RNAi)-dependent heterochromatin assembly, which was first discovered in the fission yeast, *Schizosaccharomyces pombe* (4). Factors involved in RNAi-dependent heterochromatin assembly have been comprehensively identified using biochemical, genetic, and systematic reverse genetics approaches (5, 6). In this system, small interfering RNAs (siRNAs) derived from heterochromatic regions are produced by RNAi machinery and incorporated into the RNA-induced transcriptional silencing (RITS) complex, which is composed of Ago1, Tas3, and Chp1 (4, 7). The RITS complex recruits Clr4, the sole H3K9 methyltransferase, and establishes H3K9me (8, 9). H3K9me is targeted by the HP1 proteins Swi6 and Chp2. HP1 proteins recruit the Snf2/Hdac-containing Repressor Complex (SHREC), composed of the H3K14 deacetylase Clr3, Clr1, Clr2, and the chromatin remodeling factor Mit1 to facilitate transcriptional gene silencing (10–13). Once established, heterochromatin is maintained during cell proliferation (14). However, compared with RNAi-mediated establishment, the molecular mechanism of heterochromatin maintenance is largely unknown. A few experimental systems have been developed to analyze heterochromatin maintenance in the fission yeast. One was an experimental system in which ectopic heterochromatin was established by tethering the TetR-Clr4 fusion protein to the *tetO* sequence. After the establishment of ectopic heterochromatin, Clr4 was released by adding tetracycline to analyze the subsequent maintenance of heterochromatin (15, 16). The other is a system that analyzes the process of heterochromatin maintenance by utilizing mating-type locus lacking *cis* elements such as *cenH* or *REII*, which promote heterochromatin establishment (14, 17, 18). Using these experimental systems, nuclear membrane, RNA degradation, histone turnover, cohesin, and DNA replication factors have been identified to be involved in heterochromatin maintenance (19–22). However, owing to limited experimental approaches and restricted chromosomal regions, the mechanism of heterochromatin maintenance is not fully understood. Therefore, it is desirable to develop other experimental systems that complement the aforementioned studies.

In this study, we constructed an *ade6*^+^ reporter system to study the maintenance of endogenous subtelomeric heterochromatin in the fission yeast. We demonstrated that once subtelomeric heterochromatin was established, it tended to be maintained as a metastable structure through cell proliferation. Using this system, we screened an *S. pombe* genome-wide gene deletion library for subtelomeric heterochromatin maintenance factors and successfully identified 57 factors related to DNA replication, nuclear envelope, RNA Polymerase I transcription, RNA Polymerase II transcription, structure maintenance of chromosome, ubiquitylation/sumoylation, translation, and protein folding, in addition to well-characterized heterochromatin factors. We showed that one of the factors, Mrc1^Claspin^, a mediator of the DNA replication checkpoint, maintains heterochromatin structure not only at subtelomeres, but also other heterochromatic loci such as pericentromeric and mating-type regions. In these regions, Mrc1 played a critical role in H3K9me maintenance in the absence of RNAi. In addition, we demonstrated that Mrc1 physically interacts with Clr3 and recruits SHREC to heterochromatin loci to facilitate H3K14 hypoacetylation.

## Results

### Subtelomeric chromatin as a study model of heterochromatin maintenance

Several lines of evidence have shown that RNAi and shelterin components establish heterochromatin at subtelomeres (23, 24). However, recent studies have shown that subtelomeric heterochromatin is not lost in the absence of RNAi and the shelterin component, Taz1 (20, 25). Thus, once established, subtelomeric heterochromatin appears to be maintained without establishers.

To study the details of the heterochromatin maintenance mechanism, we generated a strain (*subtel*::*ade6*^+^) harboring the *ade6*^+^ reporter gene upstream of *spac212.12* located in the subtelomeric heterochromatin on the left arm of chromosome I, in which the endogenous *ade6*^+^ was replaced with *hphMX6* (Δ*ade6*::*hphMX6*; Fig. 1A). When *ade6*^+^ is silenced by heterochromatin, cells are unable to metabolize 5-aminoimidazole ribotide, an intermediate in the adenine synthesis pathway, resulting in the formation of red colonies on low-adenine medium such as yeast extract (YE) medium (5). For example, cells with endogenous *ade6*^+^ formed white colonies (Fig. 1B), while cells with *ade6*^+^ at peri-centromeric heterochromatin (*otr1R*::*ade6*^+^) formed uniform red colonies on YE medium (Fig. 1B; (26). Once heterochromatin was compromised, colonies turned pink or white, reflecting *ade6*^+^ expression. Interestingly, cells harboring *subtel*::*ade6*^+^ formed red, pink, white, or sectored colonies on the YE medium, showing a variegation phenotype (Fig. 1B). To test epigenetic stability, red or white colonies were picked and re-streaked on the YE medium to observe colony color as they grew (Fig. 1C). Interestingly, colony colors were almost heritable across generations (red to red: 71 %, white to white: 87 %) (Fig. 1C, 1D). Importantly, these colony colors clearly reflected *ade6*^+^ mRNA expression levels (Fig. 1E). We also examined the dimethyl H3K9 (H3K9me2) level at *subtel*::*ade6*^+^ using chromatin immunoprecipitation (ChIP) analysis, and observed high H3K9me2 levels in red clones and low levels in white clones (Fig. 1F). From these observations, we concluded that once established, the subtelomeric chromatin structure is maintained and inherited as a metastable structure through cell proliferation.

**Fig. 1.**
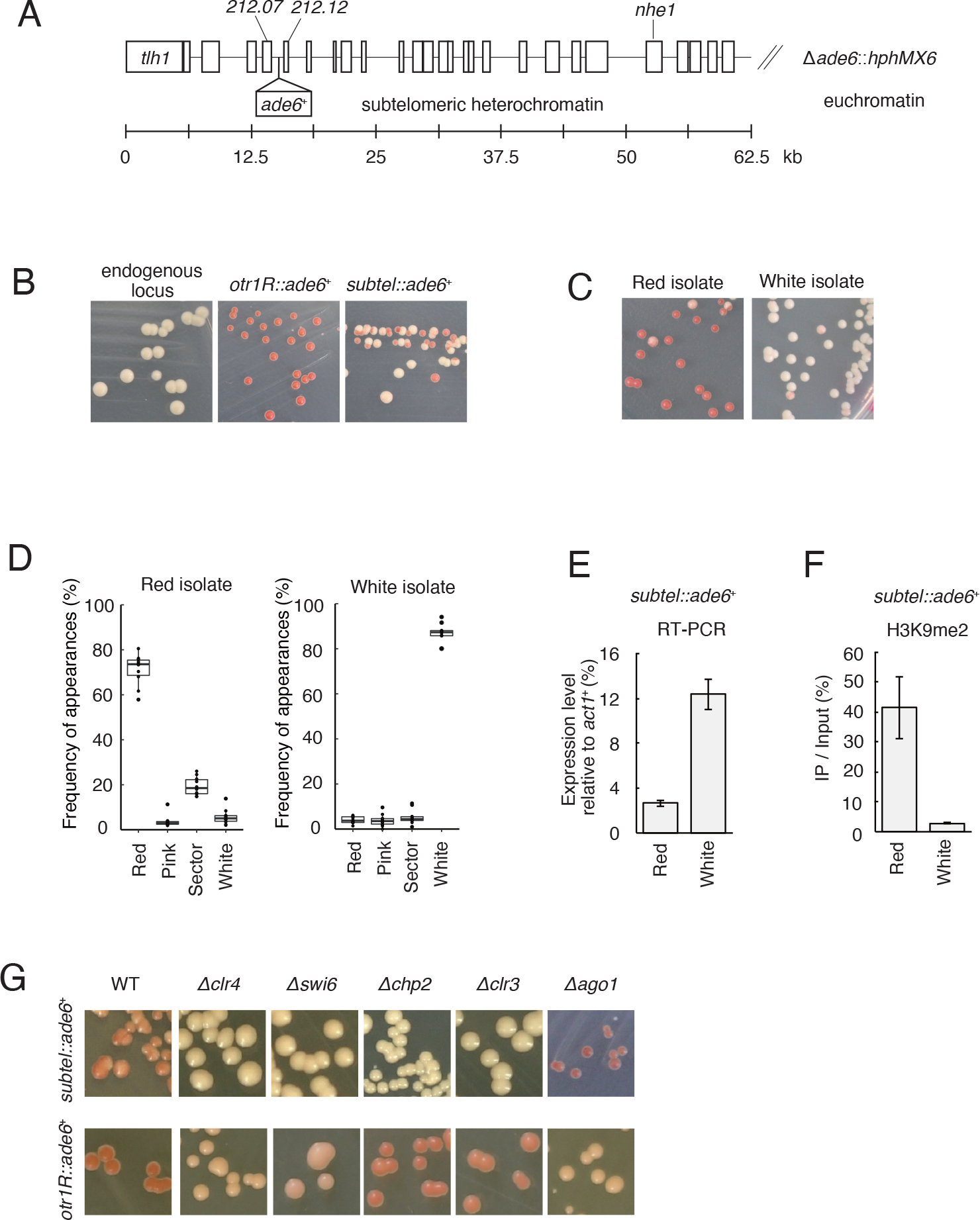
Subtelomeric chromatin as a study model of heterochromatin maintenance ***(A)*** Scheme of the subtelomeric region in the left arm of Chr I of the *subtel*::*ade6*^+^ strain. *ade6*^+^ was inserted upstream of *spac212.12* in the left arm of Chr.I. ***(B)*** Fission yeast strains carrying the *ade6*^+^ at the endogenous euchromatic locus, peri-centromere (*otr1R*::*ade6*^+^), or subtelomere (*subtel*::*ade6*^+^), were spread on the YE plates and grown for 3 d. ***(C)*** Red and white colonies derived from the *subtel*::*ade6*^+^ strain were picked and cultured in YES medium overnight, then re-spread on the YE plates and cultured for 3 d. ***(D)*** Box plot showing the quantitative results of (C). Colony color was classified into four categories: red, pink, sector, and white. The vertical axis shows the frequency of appearance of each colony; n = 10. ***(E)*** RT-qPCR analysis of *subtel*::*ade6*^+^ in the red and white isolates. The vertical axis shows the expression level of *ade6*^+^ mRNA normalized to *act1*^+^ mRNA. Error bars show standard deviation from three independent cultures. ***(F)*** ChIP analysis of H3K9me2 at *subtel*::*ade6*^+^ in the red and white isolates. The vertical axis shows the IP %. Error bars show standard deviation from three independent cultures. ***(G)*** *subtel*::*ade6*^+^ (red isolate) and *otr1R*::*ade6*^+^ strains were used as wild-type strains, and the indicated gene deletion mutants were spread on the YE plates and cultured for 3 d.

We used subtelomeric chromatin to study how heterochromatin is stably maintained and transmitted to daughter cells. First, we tested the requirement of known heterochromatin factors for the maintenance of subtelomeric heterochromatin. As expected, depletion of Clr4 and Swi6, integral components of the heterochromatin structure, resulted in the de- repression of *subtel*::*ade6*^+^ as well as *otr1R*::*ade6*^+^ (Fig. 1G). Consistent with a previous report, depletion of Ago1, an essential component of RNAi, had no effect on *subtel*::*ade6*^+^ silencing (Fig. 1G; (27). Interestingly, depletion of Chp2 or Clr3 strongly impaired *ade6*^+^ silencing at the subtelomere compared with the peri-centromere (Fig. 1G). It has been reported that Chp2 functions as a platform of SHREC (11, 13). Thus, our data suggest that subtelomeric heterochromatin maintenance and transmission require SHREC- mediated histone deacetylation.

### Systematic screening for factors involved in the maintenance of subtelomeric heterochromatin

To gain more insight into the heterochromatin maintenance mechanism, we performed a systematic screening to identify factors contributing to subtelomeric heterochromatin maintenance and inheritance. In the red isolate of the *subtel:ade6*^+^ strain, a nourseothricin resistance cassette (*natMX6*) was inserted into the *nhe1*^+^ locus located at the subtelomere on the left arm of Chr I (*nhe1-GFP*::*natMX6*) to allow selection for *subtel:ade6*^+^ (Fig. 2A). The tester strain was crossed with the *S. pombe* haploid deletion library strains (Bioneer ver 3.0) and an additional gene deletion set previously generated by the K. Gould Lab (28, 29). After sporulation, vegetative cells were killed by ethanol treatment and spores were plated onto YE plates containing G418, clonNAT, and Hygromycin B to select progenies harboring a gene deletion (Δ*geneX*::*kanMX6*), *subtel:ade6*^+^ (*nhe1- GFP*::*natMX6*) and deletion allele of endogenous *ade6*^+^ (Δ*ade6*::*hphMX6*; Fig. 2A). After growth on the selective medium, *subtel:ade6^+^*expression was assessed by colony color. Through this first screening, we isolated 79 candidates showing white or brighter pink colonies on YE plates compared to the wild type (Fig. 2A). To exclude false positives from the primary screening using the *ade6*^+^ reporter gene, we used a strain harboring *ura4*^+^ inserted into *spac212.07c* (*subtel*::*ura4*^+^) instead of *ade6^+^*(Fig. 2A). If subtelomeric heterochromatin was compromised, *ura4*^+^ would be expressed and cells would be killed on 5-FOA containing medium. Through secondary screening, 57/79 clones were found to be sensitive to 5-FOA (Fig. 2A). The *subtel*::*ade6*^+^ and *subtel*::*ura4*^+^ phenotypes of each of the 57 mutants are shown in Fig 2B and 2C, respectively. The 57 candidates were identified as subtelomeric heterochromatin maintenance factors (Table. 1). By the screening, in addition to well-characterized heterochromatin factors (*clr4*^+^, *rik1*^+^, *dos1*^+^/*raf1*^+^, *dos2*^+^/*raf2*^+^, *swi6*^+^, *chp2*^+^, *clr3*^+^, *clr1*^+^, *mit1*^+^, *ckb1*^+^, *ddb1*^+^, *cdt2*^+^, *csn1*^+^, *pob3*^+^, *swd3*^+^, *cid14*^+^, *fft3*^+^), we identified factors related to DNA replication (*mrc1*^+^, *mcl1*^+^, *rif1*^+^), nuclear envelope (*nup132*^+^, *npp106*^+^, *amo1*^+^), RNA Polymerase I transcription (*ker1*^+^, *rpa34*^+^), RNA Polymerase II transcription (*tho5*^+^, *php2*^+^, *php5*^+^*, spt3*^+^, *arp8*^+^*, cwf14*^+^), structure maintenance of chromosome (SMC) (*pds5*^+^, *nse5*^+^), ubiquitylation/sumoylation (*def1*^+^, *ubp14*^+^, *rpn10*^+^, *ubi1*^+^, *pof3*^+^, *ulp2*^+^, *rfp1*^+^), translation (*elp6*^+^, *tif310*^+^), protein folding (*ssz1*^+^, *pfd6*^+^, *btf3*^+^), and others including two unknown genes, *spbc557.02c* and *spbc1604.12*. We named *hms1*^+^ for *spbc557.02c* and *hms2*^+^ for *spbc1604.12*, respectively (<u>h</u>eterochromatin <u>m</u>aintenance at the <u>s</u>ubtelomere). Among the identified genes, *fft3*^+^, *amo1*^+^, *npp106*^+^, *mrc1*^+^, *mcl1*^+^, and *pds5*^+^ have been previously identified as factors required for the maintenance of heterochromatin in the mating-type locus or artificially induced ectopic heterochromatin (18–22, 30). Consistent with the fact that subtelomeric heterochromatin integrity does not require RNAi activity, none of the RNAi factors was identified in our screening. Overall, our screening revealed that DNA replication, RNA transcription, subnuclear structure, and protein quality/quantity control play important roles in subtelomeric heterochromatin maintenance.

**Fig. 2.**
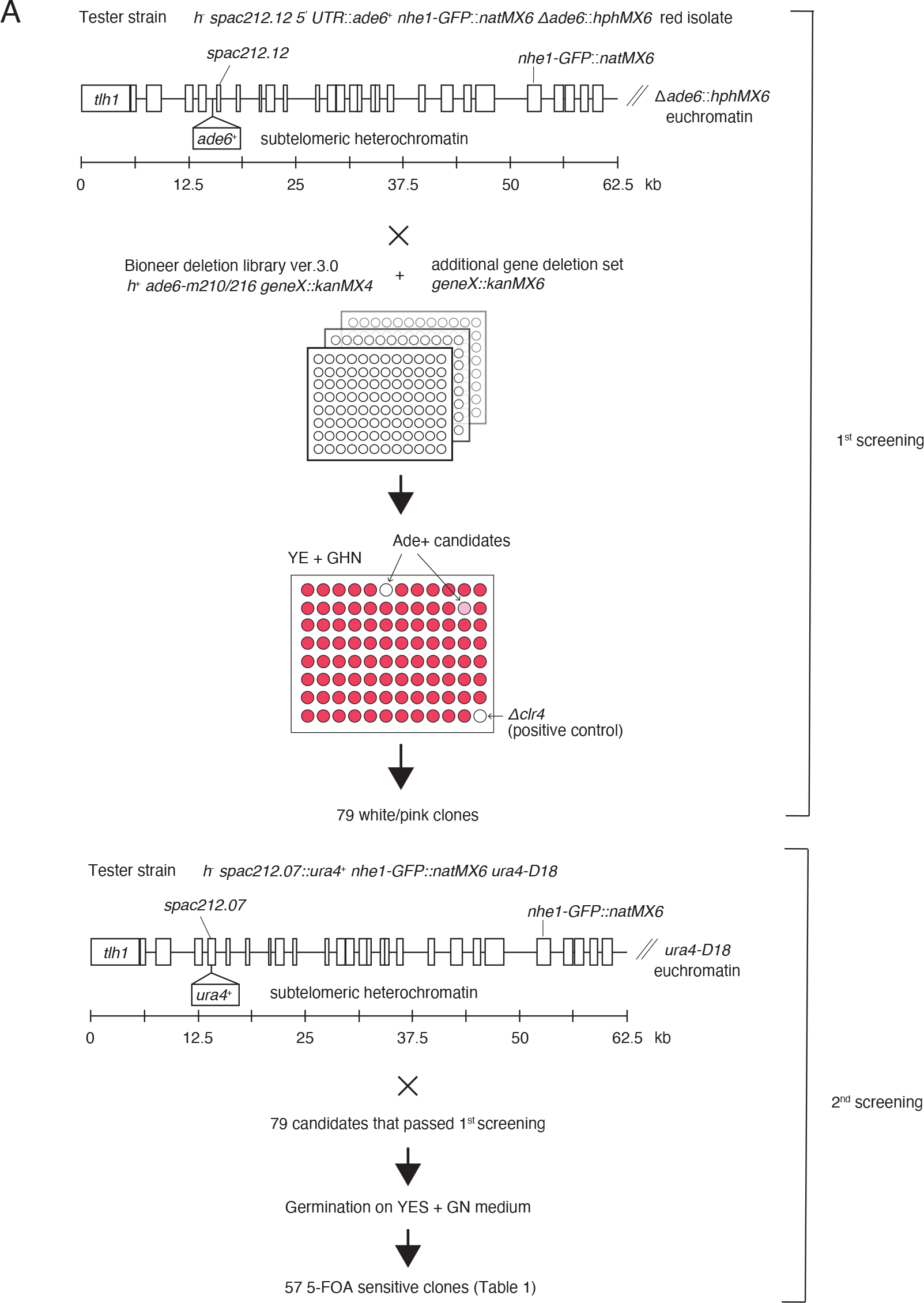
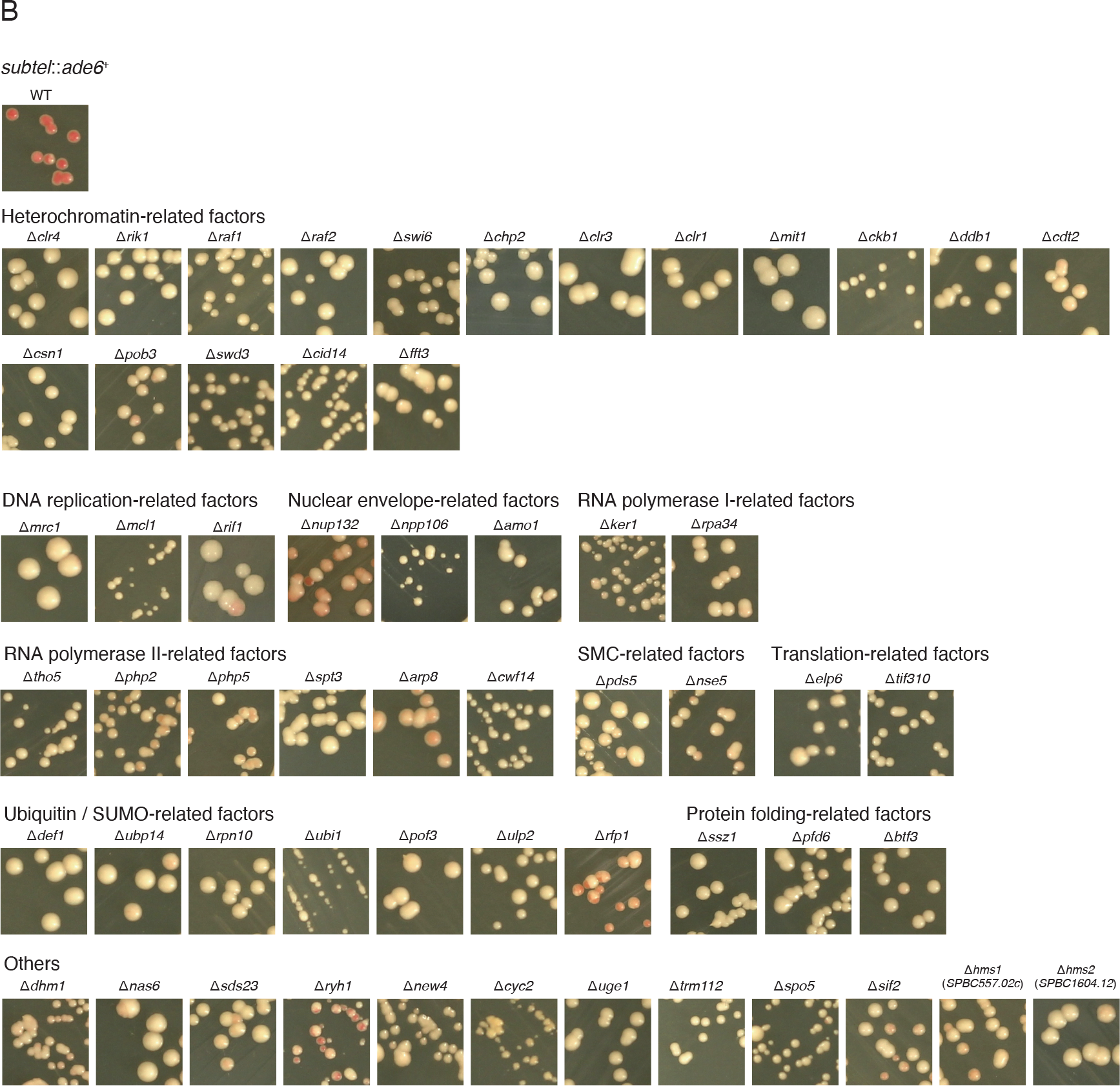
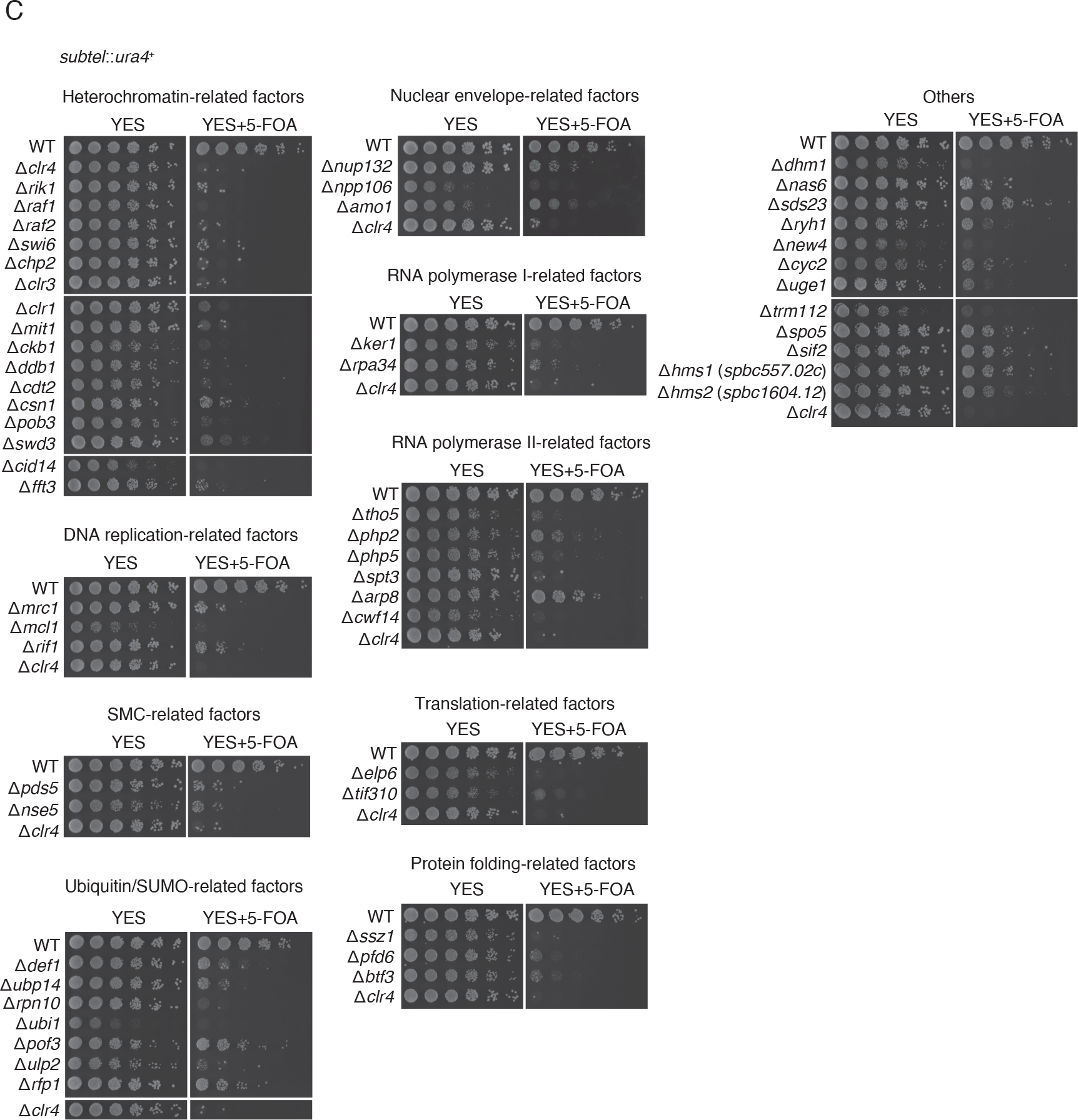
Systematic screening for factors involved in the maintenance of subtelomeric heterochromatin ***(A)*** Scheme of the systematic screening for factors involved in the maintenance of subtelomeric heterochromatin. The subtelomere on the left arm of Chr.I was genetically marked using the *natMX6* cassette inserted into the *nhe1* locus. For the primary screening, Bio5668 (*subtel*::*ade6^+^ nhe1-GFP-natMX6* Δ*ade6*::*hphMX6*) was crossed with strains of *S. pombe* haploid deletion library (Bioneer ver 3.0) and an additional gene deletion set that was previously generated (28, 29). Cells were cultured in SPAS medium for 3 d for sporulation, and were thereafter treated with 30 % ethanol for 30 min to kill vegetative cells. The progenies were spotted and germinated for 3–4 d on YE (low adenine) medium containing G418, hygromycin B, and clonNAT. 79 clones that formed whitish or pinkish colonies in comparison to the red colonies of the wild strain were selected as candidates for primary screening. In the secondary screening, KH6380 (*subtel*::*ura4*^+^ *nhe1-GFP- natMX6 ura4-D18*) was crossed with candidate strains from the primary screening. Cells were cultured in SPAS medium for 3 d for sporulation, and were thereafter treated with 30 % ethanol for 30 min to kill vegetative cells. The progenies were spread on YES medium containing G418 and clonNAT and were germinated. At least two independent G418^R^ and clonNAT^R^ clones per candidate were isolated, and their FOA sensitivities were tested. The 57 5-FOA-sensitive mutants were identified as strains that were defective in subtelomeric heterochromatin maintenance. ***(B)*** Wild-type and 57 subtelomeric heterochromatin mutants harboring *subtel*::*ade6*^+^ were spread on low adenine (YE) medium and grown for 3 d at 30 °C. ***(C)*** Five-fold serial dilutions of wild-type and 57 subtelomeric heterochromatin mutants harboring *subtel1L*::*ura4^+^* were spotted on YES and YES containing 0.1 % 5-FOA medium and grown for 3 d at 30 ℃.

**Table. 1.**
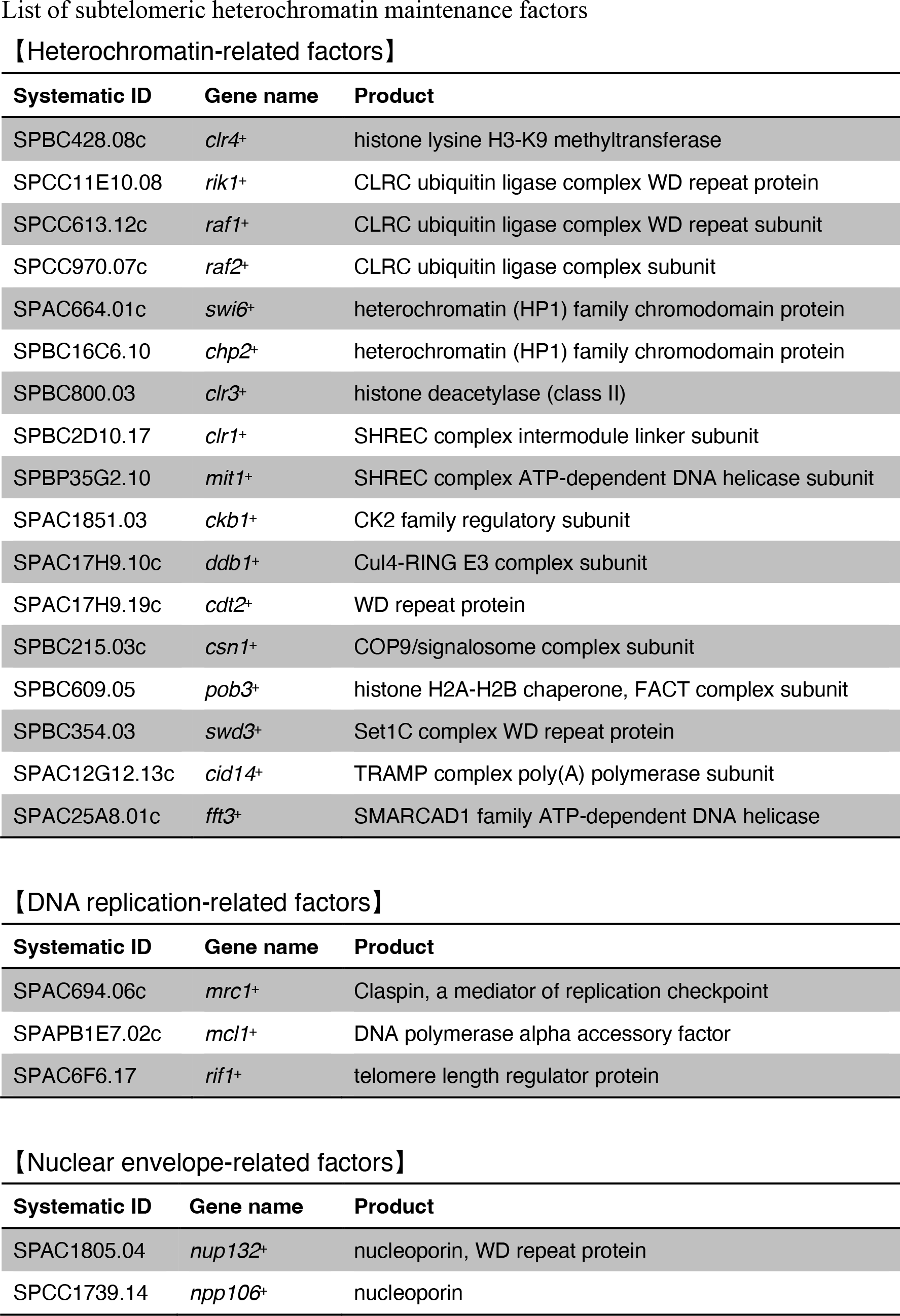
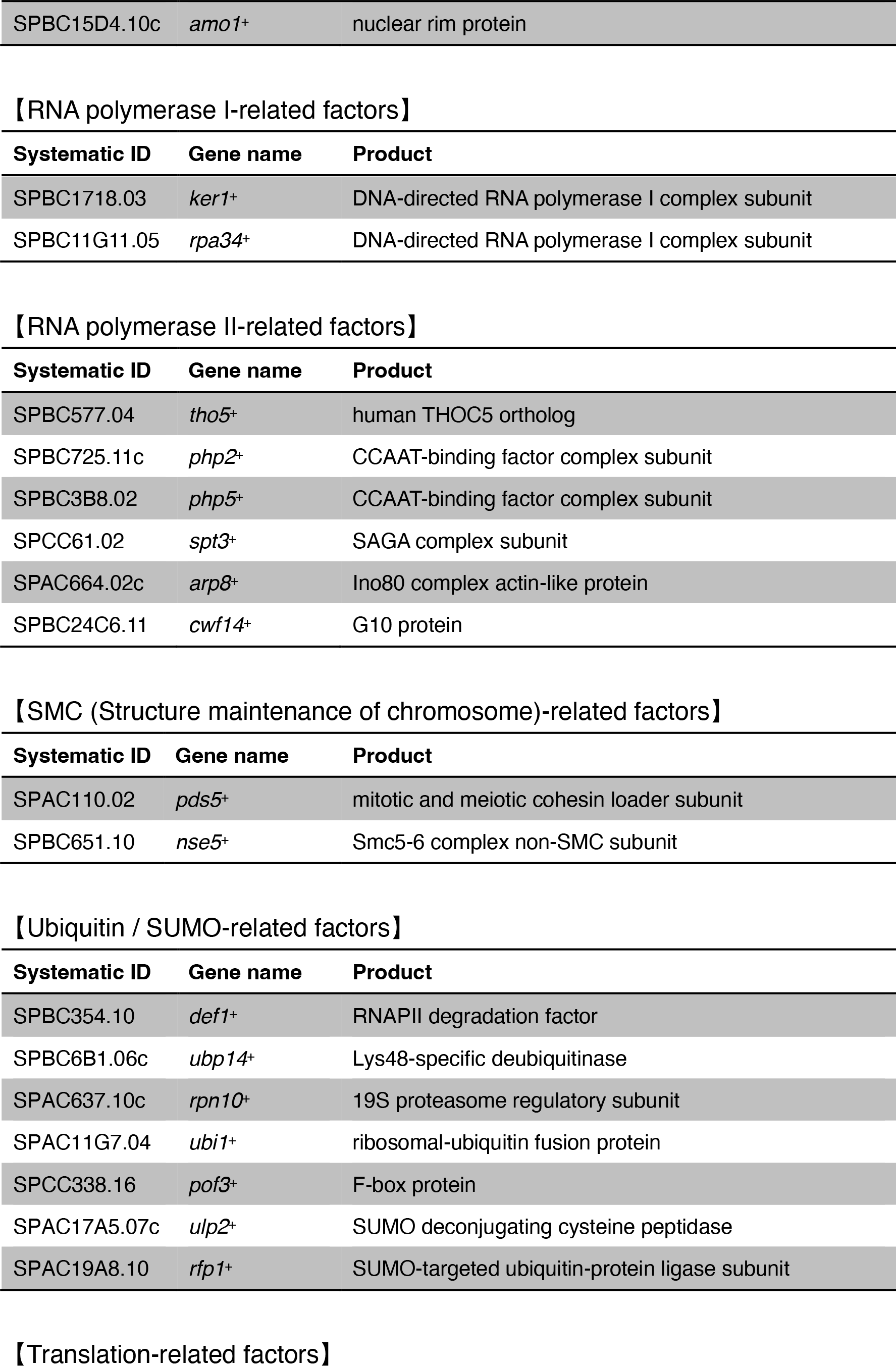
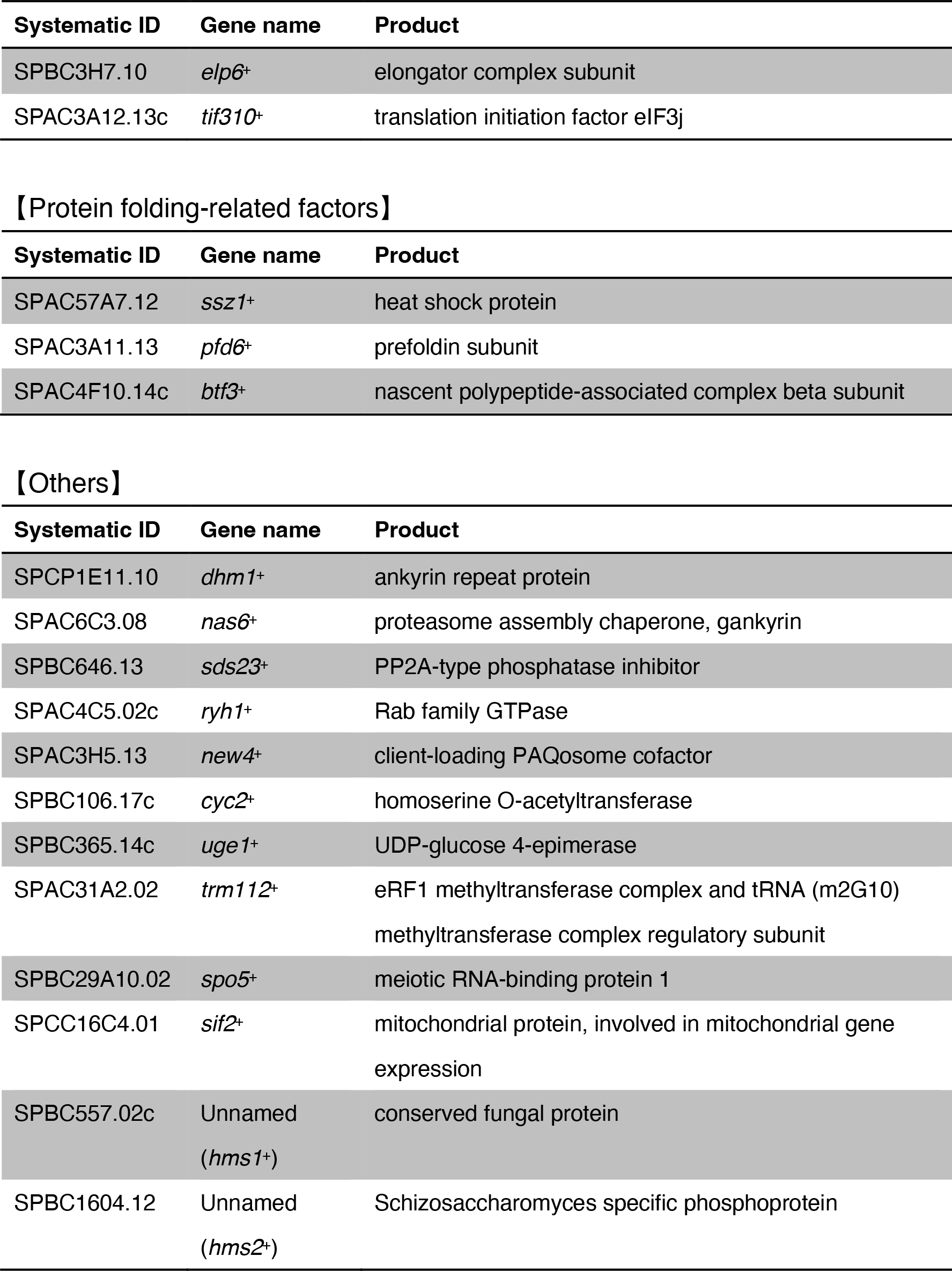
List of genes identified as factors that maintain the subtelomeric heterochromatin structure Fifty-seven factors were identified and categorized into 10 groups: heterochromatin-related, DNA replication-related, nuclear envelope-related, RNA polymerase I-related, RNA polymerase II-related, SMC-related, ubiqutin/SUMO-related, translation-related, protein folding-related factors, and others.

### Mrc1 promotes subtelomeric heterochromatin maintenance

We have previously characterized Mrc1 as a mediator of the DNA replication checkpoint (31). Since then, studies have shown that Mrc1 and its vertebrate homolog, Claspin, are involved in many aspects of S-phase regulation (32). In addition to the DNA replication-related functions, we were intrigued by Mrc1’s involvement in the maintenance of heterochromatin structure.

Mrc1 consists of several functional domains that are involved in DNA replication (Fig. 3A). The 160–317 amino acid residues on the N-terminal side form the DNA-binding domain (DBD), which facilitates binding of Mrc1 to DNA and the Swi1-Swi3 replication fork protection complex (33, 34). The central 536–674 amino acid residues contain an SQ/TQ cluster, the phosphorylation site of Rad3 and Tel1, required for the activation of the replication checkpoint (35). The 782–879 amino acid residues are named the Hsk1- bypass segment (HBS), which represses weak-early firing origins (36). However, the function of the C-terminal 880–1019 amino acid region is unknown.

**Fig. 3.**
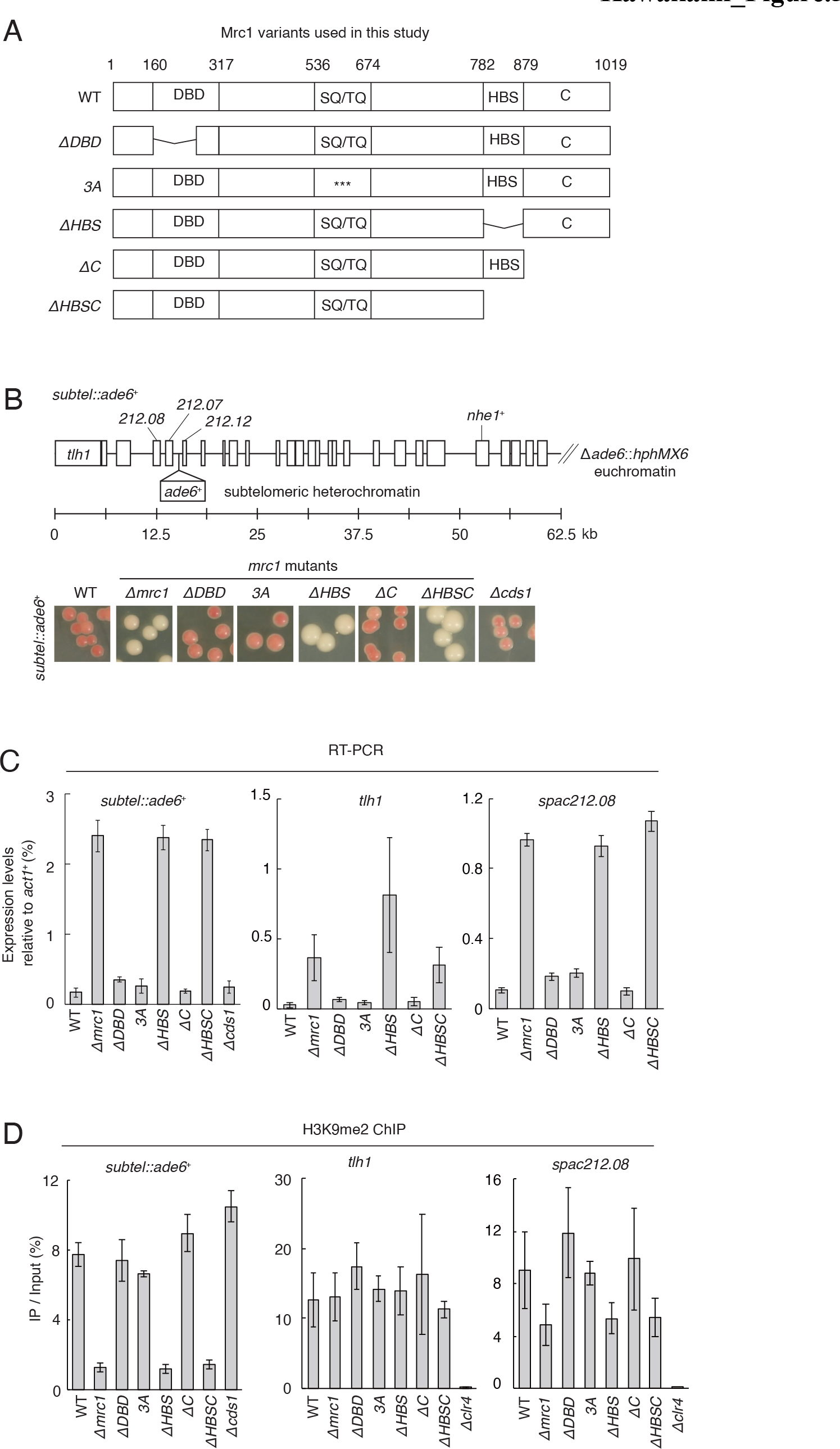
Mrc1 promotes subtelomeric heterochromatin maintenance ***(A***) Schematic diagram of the domain structure and variants of Mrc1 protein. Wild-type, deletion of amino acids from 160-284 (Δ*DBD*), 782-879 (Δ*HBS*), 880-1019 (Δ*C*), 782- 1019 (Δ*HBSC*), or mutation S604A T645A T653A (*3A*) in SQ/TQ of Mrc1 proteins. ***(B)*** Scheme of subtelomeric region in the left arm of Chr I in *subtel*::*ade6*^+^ strain. *ade6*^+^ was inserted into the upstream of *spac212.12* (*subtel*::*ade6*^+^) (upper). Indicated strains harboring *subtel*::*ade6*^+^ were streaked on YE plates and incubated for 3 d (lower). ***(C)*** RT-qPCR analysis of *subtel*::*ade6*^+^, *tlh1*, and *spac212.08* in indicated strains. The vertical axis shows mRNA expression levels relative to *act1*^+^ mRNA. ***(D)*** ChIP analysis of H3K9me2 at *subtel*::*ade6*^+^, *tlh1*, and *spac212.08* in indicated strains. The vertical axis in ChIP analysis shows IP %. The chromosomal positions of *tlh1* and *spac212.08* were indicated in *(B)*. Error bars in *(C)* and *(D)* show the standard deviation from three independent cultures.

To identify the functional domain of Mrc1 for heterochromatin maintenance at the subtelomere, we generated each domain mutant on a *subtel*::*ade6^+^* genetic background (Fig. 3A). Δ*HBS* and Δ*HBSC* cells formed white colonies similar to the Δ*mrc1* cells, while the Δ*DBD*, *3A* (S604A/T645A/T653A, an SQ/TQ mutant specifically defective in checkpoint function), or Δ*C* cells formed red colonies similar to the wild-type on the YE medium (Fig. 3B). These results clearly showed that HBS was responsible for subtelomeric gene silencing, whereas neither DNA binding, fork protection, nor replication checkpoint function was required. Consistent with this result, our screening did not identify Swi1 or Swi3 (Table.1). In addition, depletion of *cds1^+^*, a replication checkpoint kinase had no defect on *subtel*::*ade6^+^* silencing (Fig. 3B). These results were confirmed using the *ura4*^+^ marker gene instead of *ade6*^+^ (*subtel*::*ura4*^+^). Δ*mrc1*, Δ*HBS*, or Δ*HBSC* exhibited strong sensitivity to 5-FOA, suggesting *ura4*^+^ de-repression (Fig. S1A). To confirm the results of the spot assay, we measured *ade6*^+^ mRNA expression levels by RT-qPCR (Fig. 3C). The level of mRNA derived from *subtel*::*ade6*^+^ was elevated in Δ*mrc1,* Δ*HBS*, and Δ*HBSC* cells, but not in *3A*, Δ*DBD*, and Δ*C* cells (Fig. 3C). We also measured the expression levels of endogenous the subtelomeric genes, *tlh1* and *spac212.08*, and obtained similar results to those of *subtel*::*ade6*^+^ (Fig. 3C). Next, to analyze heterochromatic histone modification in *mrc1* mutants, we performed ChIP analysis of H3K9me2 (Fig. 3D). At *subtel*::*ade6*^+^ and *spac212.08*, H3K9me2 levels were reduced in Δ*mrc1*, Δ*HBS,* and Δ*HBSC* cells, but did not change in *3A*, Δ*DBD*, and Δ*C* cells (Fig. 3D). Similar results were observed for *subtel*::*ura4*^+^ (Fig. S1B). However, even in Δ*mrc1* cells, H3K9me2 levels did not change at *tlh1,* despite an increase in mRNA levels (Fig. 3C and 3D). Overall, we conclude that Mrc1 promotes gene silencing and H3K9me2 in subtelomeric heterochromatin. We also mention that the decrease in H3K9 methylation cannot generally explain the silencing defect of *mrc1* mutant cells (e.g., *tlh1* locus; Fig. 3C and 3D).

### Mrc1 is required for the maintenance of heterochromatin independently of RNAi

Next, we tested the requirement of Mrc1 for the maintenance of other heterochromatic loci, such as the peri-centromeres and mating-type regions. To monitor the gene expression at the peri-centromeres, we used a strain harboring *otr1R*::*ade6^+^*(Fig. 4A). Δ*mrc1* cells formed dark red colonies on the YE plate, as observed in wild-type cells, indicating that Mrc1 is not required for gene silencing at the peri-centromeres (Fig. 4A). Alternatively, Δ*ago1* cells formed brighter pink colonies, similar to Δ*clr4* cells, reflecting the importance of the RNAi machinery for pericentromeric heterochromatin (Fig. 1G and 4A). Consistent with the result of colony color assay, the levels of H3K9me2 on peri- centromeric *dg* repeats in *mrc1* mutants showed the same level as those of wild-type, but was greatly reduced in Δ*ago1* cells (Fig. 4B). These results suggest that Mrc1 is not required for RNAi-mediated heterochromatin assembly at peri-centromeres.

**Fig. 4.**
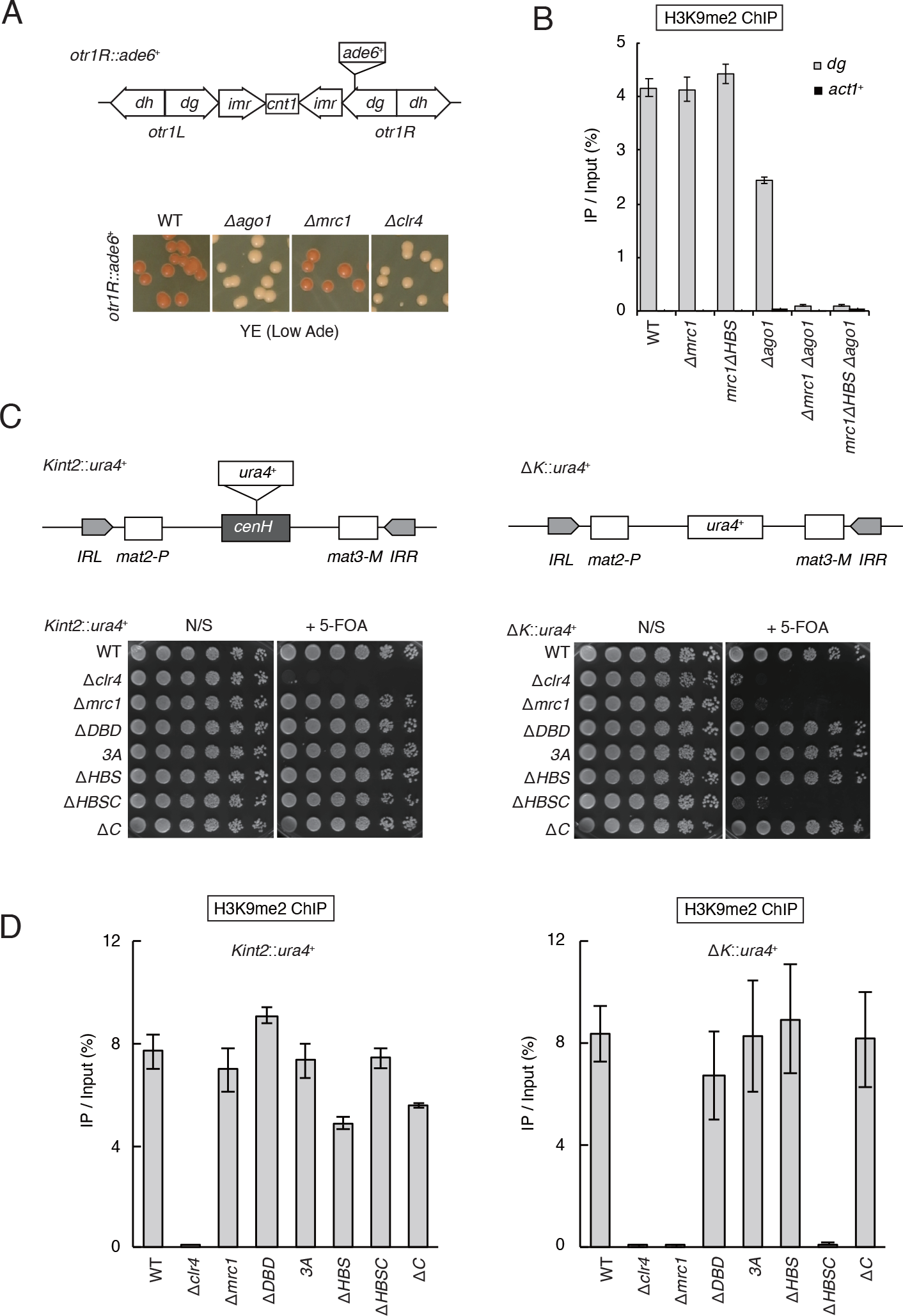
Mrc1 is required for heterochromatin maintenance independently of RNAi ***(A***) Scheme of the centromeric region of Chr I with *ade6*^+^ insertion (upper panel). Five- fold serial dilutions of the indicated genotype cells harboring *otr1R*::*ade6^+^*were spotted onto YES and YE and incubated for 3 d at 30 °C (lower). ***(B)*** ChIP analysis of H3K9me2 in pericentromeric *dg* repeats from the indicated strains. The *act1^+^* locus was used as the control for euchromatin. The vertical axis shows the IP %. ***(C)*** Five-fold serial dilutions of indicated genotype cells harboring *kint2*::*ura4^+^* (left) or Δ*K*::*ura4*^+^ (right) were spotted onto YES (N/S) and YES + 0.1 % 5-FOA (+ 5-FOA) and incubated for 3 d. ***(D)*** ChIP analysis of H3K9me2 at *kint2*::*ura4^+^* (left) or Δ*K*::*ura4*^+^ (right) in indicated strains. The vertical axes indicate IP %. Error bars in *(B)* and *(D)* show standard deviations from three independent cultures.

Although RNAi promotes the establishment of heterochromatin at the peri-centromeres, in the absence of RNAi, some portion of H3K9me2 remained (Fig. 4B). To test the requirement of Mrc1 for H3K9me2 maintenance in the absence of RNAi, we examined H3K9me2 levels in *mrc1* and *ago1* double-mutant cells (Fig. 4B). As a result, Δ*mrc1* Δ*ago1* and *mrc1*Δ*HBS* Δ*ago1* cells failed to maintain the H3K9me2 at the peri- centromere (Fig. 4B), indicating that Mrc1 is essential for the maintenance of peri- centromeric heterochromatin in the absence of RNAi.

Next, we tested the requirement of Mrc1 for heterochromatin maintenance in the mating-type locus. In the mating-type locus, the flanking region between *mat2-P* and *mat3-M* has the *cenH* sequence that is homologous to peri-centromeric repeats, promoting RNAi-mediated heterochromatin establishment (14, 37). Unlike the insertion of *ura4*^+^ into the *cenH* region (*kint2*::*ura4*^+^), the replacement of *cenH* with *ura4*^+^ (Δ*K*::*ura4*^+^) invalidates the *de novo* heterochromatin assembly in the mating-type locus. In the absence of the *cenH* element, heterochromatin becomes unstable and produces *ura4*-on and *ura4*-off clones (17). However, once the *ura4-*off state is established, the silent chromatin state tends to be inherited in *cis* (17). Depletion of *mrc1* caused silencing defects in the Δ*K*::*ura4^+^* (*ura4*-off) background, whereas no silencing defect was observed in the *Kint2*::*ura4*^+^ background (Fig. 4C). Consistent with this, our ChIP analysis revealed that depletion of Mrc1 caused a dramatic reduction in the H3K9me2 levels of Δ*K*::*ura4^+^*, but caused a marginal effect in the *Kint2*::*ura4^+^* genetic background (Fig. 4D). These results clearly show that Δ*mrc1* cells are proficient in heterochromatin structure assembly when RNAi-mediated de novo assembly mechanisms are active, but fail to maintain heterochromatin in the absence of RNAi. Interestingly, HBS depletion did not cause heterochromatin defects in the Δ*K*::*ura4^+^* background, whereas HBSC depletion significantly compromised gene silencing and H3K9me2 (Fig. 4C and 4D). Thus, in addition to HBS, the C-terminal region of Mrc1 is important for the maintenance of heterochromatin, specifically in the mating-type locus.

Through our analysis of the peri-centromere and mating-type regions, we concluded that Mrc1 is generally required for heterochromatin maintenance in the absence of an RNAi-dependent heterochromatin establishment pathway.

### Mrc1 maintains the hypoacetylation state at H3K14 by recruiting SHREC

Our analysis showed that the mutation of Mrc1 leads to de-repression of *tlh1* mRNA without the loss of H3K9me2 (Fig. 3C and 3D). Heterochromatic gene silencing is ensured not only by H3K9me but also by the hypoacetylation state at H3K14. Since SHREC, an H3K14 deacetylation complex, is required for gene silencing at subtelomeric heterochromatin like Mrc1 (Fig. 1G, 3B), we hypothesized that Mrc1 may be involved in the hypoacetylation state of H3K14. To test this, we performed ChIP analysis of H3K14ac in *mrc1* mutants. The results showed that the level of H3K14ac was elevated in the subtelomeric and pericentromeric heterochromatin in the Δ*mrc1* and Δ*HBS* mutants compared to that in the wild-type (Fig. 5A). This result suggests that Mrc1 is important for the hypoacetylation of H3K14. As H3K14 is deacetylated by SHREC, we considered the possibility that Mrc1 is involved in the SHREC-dependent pathway. It had been reported that two HP1 proteins, Swi6 and Chp2, promote a hypoacetylation state at H3K14 by recruiting SHREC (11, 12). We therefore analyzed the localization of these two HP1 proteins and Mit1, a subunit of SHREC, on heterochromatin in *mrc1* mutants. The results of ChIP analysis showed that occupancies of Swi6 and Chp2 were slightly decreased at *spac212.08* (Fig. 5B and 5C). Importantly, Swi6 and Chp2 were still localized at *tlh1* and *dg*, whereas their H3K9me2 levels were not affected in *mrc1* mutants (Fig. 3D, 4B, 5B, and 5C). This result suggested that the HP1 occupancy reflected the residual levels of H3K9me at each locus in *mrc1* mutants (Fig 3D, 4B, 5B, and 5C). Thus, Mrc1 does not directly regulate the localization of HP1 proteins that are the SHREC platform. In contrast, Mit1 localization at the subtelomeric and peri-centromeric heterochromatin was abolished in *mrc1* mutants (Fig. 5D). These results suggest that Mrc1 maintains a hypoacetylation state at H3K14 by recruiting SHREC to HP1 proteins. We previously reported that Mrc1 is predominantly expressed during the S phase (31).

**Fig. 5.**
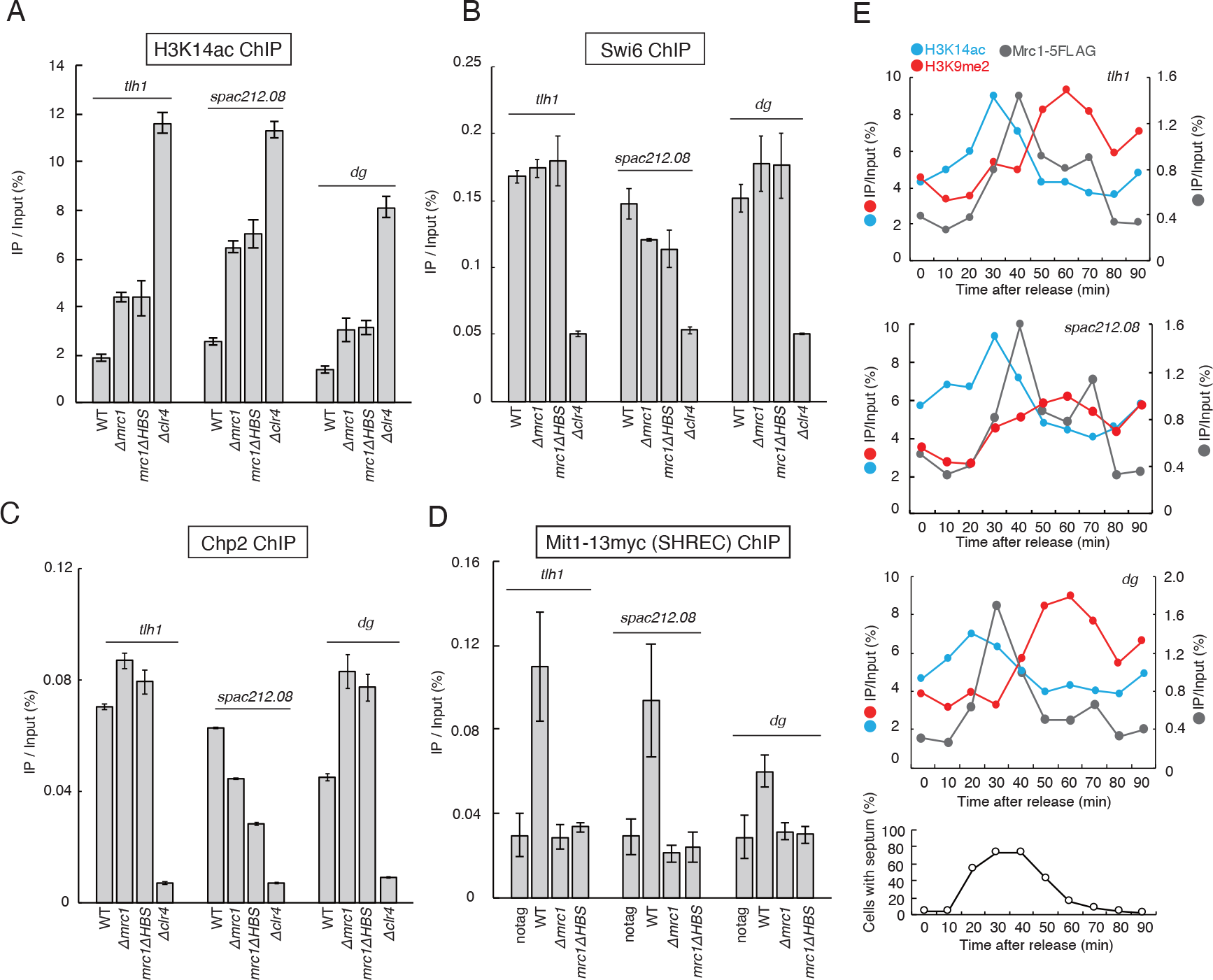
Mrc1 maintains the hypoacetylation state of H3K14 by recruiting SHREC. ChIP analysis of H3K14ac ***(A)***, Swi6 ***(B)***, Chp2 ***(C)***, and Mit1-13myc ***(D)*** in subtelomeric (*tlh1*, *spac212.08*) and pericentromeric (*dg*) regions in the indicated strains. The vertical axes indicate IP %. Error bars in *(A)*, *(B)*, *(C)*, and *(D)* show standard deviations from three independent cultures. ***(E)*** ChIP analysis of H3K14ac (blue), H3K9me2 (red), and Mrc1-5FLAG (gray) at *tlh1*, *spac212.08,* and *dg* during the cell cycle in *nda3-KM311* background. The main vertical axes show IP % for H3K14ac and H3K9me2. The secondary vertical axes show IP % for Mrc1-5FLAG. The horizontal axes show the time (min) after release. The bottom panel shows septation index. The cells within septa were counted and plotted; at least 150 cells were counted for each time point.

In addition, Mrc1 is involved in many aspects of S-phase regulation (32). These reports prompted us to analyze Mrc1 localization and histone modifications in heterochromatin throughout the cell cycle. We used *nda3-KM311* cold-sensitive mutant cells to arrest the cell cycle at mitotic prophase and grow synchronously (38, 39). Cell cycle progression was monitored by the septation index (the percentage of cells with septa), which peaked during the S phase (Fig. 5E). H3K14ac levels increased markedly in the early/mid S phase and rapidly decreased at late S/G2 in the subtelomeric (*tlh1* and *spac212.08*) and pericentromeric (*dg*) heterochromatin (Fig. 5E). In contrast to H3K14ac, H3K9me2 levels were low in the S phase and rapidly increased at the S/G2 transition (Fig. 5E). This result is consistent with previous reports that heterochromatin is derepressed during the S phase to allow transcription by RNA polymerase II (40). Interestingly, the occupancy of Mrc1 showed a clear peak at the H3K14ac/H3K9me2 transition phase in the subtelomeric and pericentromeric heterochromatin (Fig. 5E). These results suggest that Mrc1 may reduce H3K14ac in the late S phase by recruiting SHREC.

### Mrc1 physically interacts with Clr3 via Mrc1’s HBS containing C-terminal region

Based on the result showing that Mrc1 contributes to SHREC recruitment (Fig. 5D), we examined the physical interactions between Mrc1 and SHREC using a yeast two-hybrid assay (Y2H). Full-length Mrc1 (Mrc1-F) was expressed as a fusion protein that contained a DNA-binding domain of Gal4 (Gal4-BD; Fig. 6A), while each component of SHREC, Clr1, Clr2, Clr3, or Mit1 was expressed as a fusion protein with a transcriptional activation domain of Gal4 (Gal4-AD). If the two proteins bind to each other, transcriptional activation of the *HIS3* reporter gene results in histidine biosynthesis and yeast growth on histidine-deficient media. In our Y2H experiments, co-expression of Mrc1-F and Clr3 resulted in a positive Y2H interaction, suggesting a physical interaction between Mrc1 and Clr3 (Fig. 6B). To determine the region in Mrc1 that is sufficient for interaction with Clr3, we generated three Mrc1 derivatives, Mrc1-N (1–503 a.a.), Mrc1- M (248–756 a.a.), and Mrc1-C (500–1019 a.a.) fused to Gal4-BD and performed Y2H experiments again (Fig. 6A and C). Mrc1-C was sufficient for interaction with Clr3 (Fig. 6C). Although Mrc1-M and Mrc1-C share 500–756 aa, Mrc1-M did not interact with Clr3 (Fig. 6C), suggesting that the 757–1019 aa C-terminal region containing HBS is important for the interaction with Clr3. Taken together, we conclude that Mrc1 physically interacts with Clr3 via Mrc1’s HBS containing C-terminal region.

**Fig. 6.**
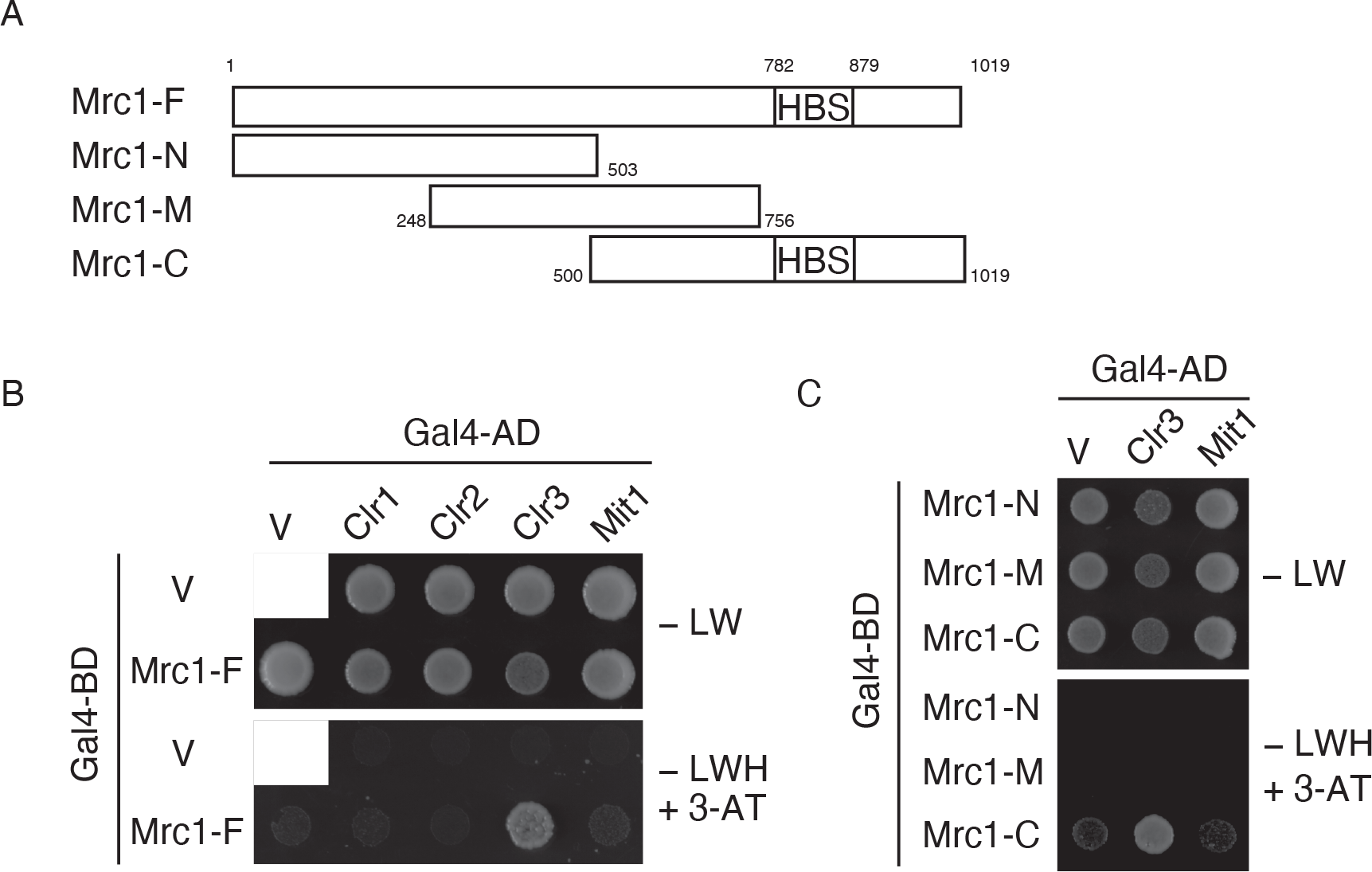
Mrc1 physically interacts with Clr3 via Mrc1’s HBS containing C-terminal region ***(A)*** Schematic diagram of each Mrc1 fragment used in yeast two-hybrid assay (Y2H). The full length (Mrc1-F), 1–503 a.a. N terminal region (Mrc1-N), 248–756 a.a. middle region (Mrc1-M), and 500–1019 a.a. C-terminal regions (Mrc1-C) were fused with the Gal4 DNA binding domain (Gal4-BD). ***(B)*** Y2H assay between Mrc1-F and SHREC components. Full length Clr1, Clr2, Clr3, or Mit1 were fused with Gal4-AD and co-expressed with Mrc1-F-Gal4-BD in a *S. cerevisiae* AH109 strain, which harbors the *HIS3* reporter gene. V stands for empty vector. Cells were spotted onto SC-LEU-TRP (-LW) or SC-LEU-TRP-HIS+ 20 mM 3-Amino-1,2,4-triazole (3-AT) and incubated at 30 ℃. ***(C)*** Y2H assay between Mrc1 fragments (Mrc1-N, -M, or -C) and Clr3. Empty vector and Mit1 served as negative controls. Transformants were spotted onto SC-LEU-TRP (-LW) or SC-LEU-TRP-HIS (-LWH) + 100 mM 3-AT and incubated at 30 ℃.

### Depletion of Mst2, a H3K14 acetyltransferase, restores heterochromatin defects in *mrc1* mutants

If Mrc1 facilitates the hypoacetylation state of H3K14 to maintain the heterochromatin structure, it is possible that artificial reduction of H3K14ac levels rescues the silencing defect in *mrc1* mutants. To test this possibility, we depleted Mst2, an H3K14-specific acetyltransferase, in *mrc1* mutants. Supporting our hypothesis, Mst2 depletion significantly suppressed the de-repression of *subtel*::*ura4*^+^ in Δ*mrc1* and *mrc1*Δ*HBS* mutant cells (Fig. 7A). Moreover, Mst2 depletion restored H3K9me2 at subtelomeres in Δ*mrc1* or Δ*HBS* mutants to almost the same levels as that of the wild-type (Fig. 7B). In the mating-type locus, Mst2 depletion suppressed the de-repression of Δ*K*::*ura4*^+^ in the Δ*mrc1* or *mrc1*Δ*HBSC* mutants (Fig. 7C). Mst2 depletion also restored H3K9me2 levels in Δ*K*::*ura4*^+^ in Δ*mrc1* or Δ*HBSC* mutants (Fig. 7D). Collectively, these findings demonstrate that suppressing H3K14ac is sufficient to explain Mrc1’s function in the maintenance of heterochromatin.

**Fig. 7.**
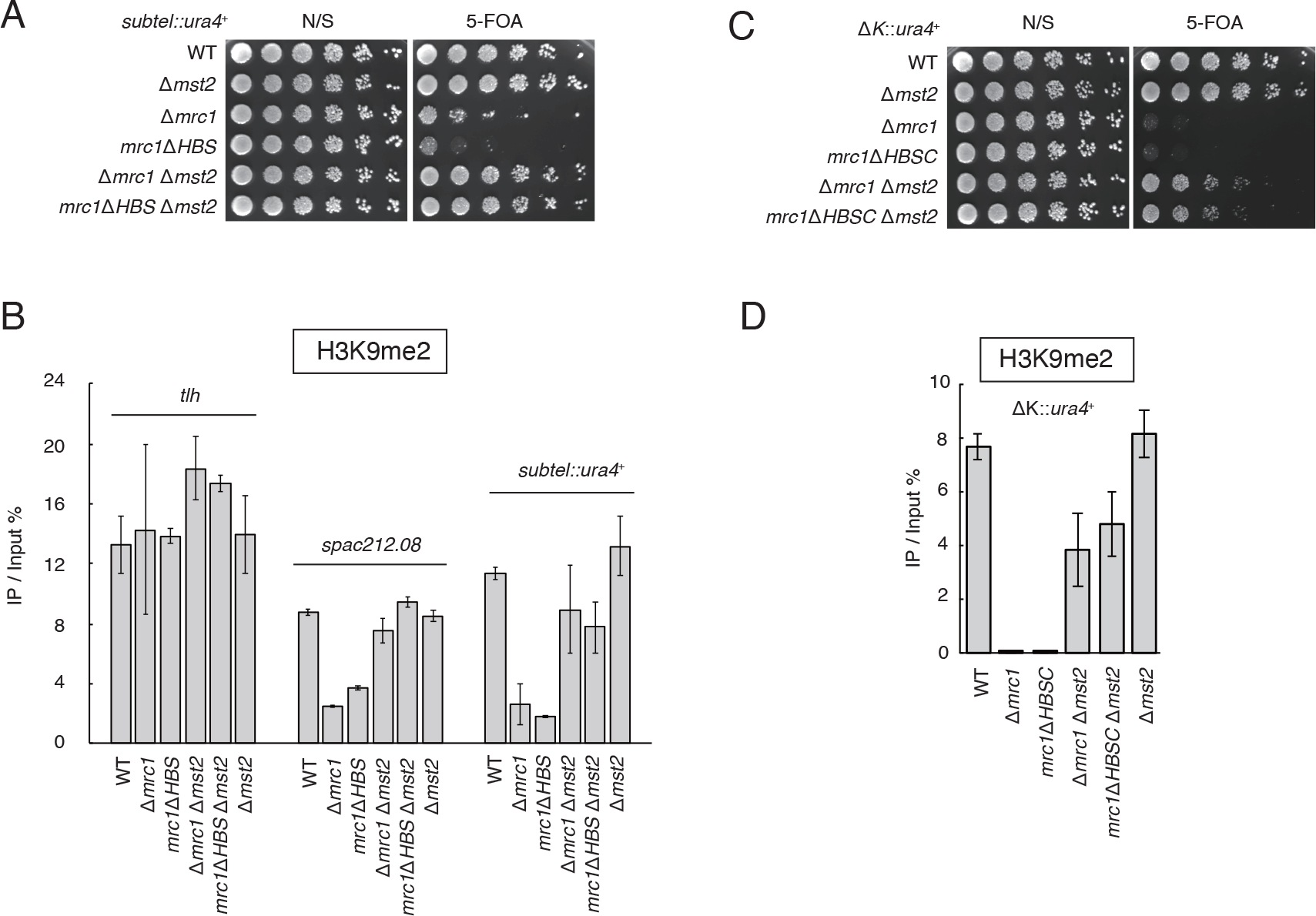
Depletion of Mst2, a H3K14 acetyltransferase, restores heterochromatin defects in *mrc1* mutants ***(A)*** Five-fold serial dilutions of indicated genotype cells harboring *subtel*::*ura4^+^* were spotted onto YES (N/S) and YES + 0.1 % 5-FOA (5-FOA) plates and incubated for 3 d at 30 ℃. ***(B)*** ChIP analysis of H3K9me2 in the subtelomeric regions of the indicated strains. The vertical axis shows the IP %. ***(C)*** Five-fold serial dilutions of indicated genotype cells harboring Δ*K*::*ura4^+^* were spotted onto YES (N/S) and YES + 0.1 % 5- FOA (5-FOA) plates and incubated for 3 d at 30 ℃. ***(D)*** ChIP analysis of H3K9me2 in Δ*K*::*ura4^+^* in the indicated strains. The vertical axis shows IP %. Error bars in *(B)* and *(D)* show the standard deviations from three independent cultures.SHREC

## Discussion

In this study, we constructed a novel evaluation system for heterochromatin maintenance in subtelomeres (Fig. 1). We showed that SHREC-mediated histone deacetylation has more prominent effects on heterochromatin integrity at subtelomeres than at peri- centromeres (Fig. 1G). In addition, our screening revealed that the heterochromatin maintenance process is associated with various nuclear functions (Table. 1 and Fig. 2). This is the first report of a systematic screening for heterochromatin maintenance factors in endogenous subtelomeric heterochromatin. We showed that one of the identified factors, Mrc1, is linked to SHREC to maintain the heterochromatin structure not only at the subtelomere but also at the pericentromeric regions and mating-type locus (Fig. 4). Mrc1 associates with heterochromatin during the S phase, suggesting that Mrc1 may coordinate DNA replication and heterochromatin maintenance (Fig. 5E). The results of Mrc1 analysis suggest that subtelomeric heterochromatin maintenance factors possibly influence heterochromatin stability more generally, not just at the subtelomeres (Fig. 4). However, it remains unclear how individual genes promote heterochromatin maintenance. Elucidation of the molecular functions of individual genes is necessary to gain a complete understanding of heterochromatin maintenance mechanisms. The details of these findings are discussed below.

### Epigenetic inheritance of subtelomeric chromatin structure

In this study, we showed that the gene expression state was relatively unstable in fission yeast subtelomeres (Fig. 1B). This result suggests that the mechanisms that positively and negatively regulate heterochromatin are antagonistic at the subtelomeres (Fig. 1B). However, once established, the expression states became metastable throughout the generations (Fig. 1C and 1D). Interestingly, both ON and OFF states of gene expression were maintained almost equally (Fig. 1C and 1D). Variable and metastable phenotypes of subtelomeric chromatin are similar to the phenotype observed in the Δ*K* strain in which the *de novo* heterochromatin assembly at the mating-type locus is invalidated (17). Thus, at subtelomeres, the heterochromatin establishment pathway may be inefficient, similar to the Δ*K* mating-type locus.

At the subtelomere, RNAi and Taz1 redundantly function in heterochromatin establishment (23). Importantly, a previous report showed that double depletion of RNAi protein and Taz1 did not affect H3K9me levels at subtelomeres (25). Thus, it seems that subtelomeric heterochromatin can be maintained without RNAi or Taz1 after establishment. Unlike centromeres, where RNAi-dependent recruitment of Clr4 occurs actively, self-maintenance mechanisms seem to play a more central role in heterochromatin integrity at subtelomeres. Our data revealed that Clr3-mediated histone H3 deacetylation is essential for the maintenance of subtelomeric but not peri-centromeric heterochromatin (Fig. 1G). This is consistent with a previous report that SHREC and Chp2 are required for the maintenance of ectopically induced heterochromatin (19). In addition to Clr3, our screening identified many factors related to histone modification, DNA replication, nuclear envelope, RNA transcription, SMC, ubiquitin/SUMO, translation, and protein folding (Table. 1). In cells, DNA replication and RNA polymerase II transcription processes that require the turnover of histones and/or nucleosomes are potential risks to the stable maintenance of histone modifications (41, 42). Therefore, it appears that DNA replication and RNA polymerase II transcription machineries are equipped with a mechanism to stably pass histone modifications to the next generation. Interestingly, we identified two RNA polymerase I-specific subunits – Ker1 and Rpa34 (Table. 1). RNA polymerase I catalyzes rRNA transcription in the nucleolus. Since telomeres are clustered at the nuclear periphery near the nucleolus in fission yeast (43, 44), it is possible that subtelomeric heterochromatin maintenance requires nucleolar attachment or nucleolar proteins such as RNA polymerase I. Identification of the nuclear envelope and SMC-related factors suggested that the stable maintenance and transmission of the epigenetic state is possibly linked to subnuclear 3D chromatin organization (18, 20, 22, 45, 46). The finding of several ubiquitination, sumoylation, translation, and protein folding related factors indicated that fine-tuning of protein quantity, quality, or function is also essential for the maintenance of heterochromatin. Mrc1, Mcl1, and Npp106, which we identified in this screen, were also previously identified as factors that function specifically in the maintenance of ectopically generated heterochromatin by tethering the TetR-Clr4 system (19, 20). Other factors, including Pds5, Amo1, and Fft3, which we identified in this screen, have also been reported to promote the epigenetic inheritance of heterochromatin in the mating-type locus (18, 21, 22). Thus, subtelomeres are a sophisticated study model for the epigenetic inheritance of heterochromatin, along with the above artificial systems or other chromosomal regions.

### SHREC recruitment

Our results showed that SHREC was essential for the maintenance of subtelomeric heterochromatin (Fig. 1G). Because SHREC is critical for H3K14 deacetylation and heterochromatic gene silencing (10), several pathways ensure the recruitment of SHREC to HP1. The phosphorylation of Swi6 and Chp2 by CKII promotes SHREC recruitment (47). Another report showed that the nuclear membrane protein, Lem2, facilitates SHREC recruitment by regulating the peripheral nuclear localization of heterochromatin (25). These reports suggest that SHREC recruitment is regulated temporally and spatially in the nucleus. In addition, we showed that Mrc1 localizes to heterochromatin during the S phase and is required for the chromatin localization of SHREC (Fig. 5D and 5E), suggesting the possibility that Mrc1 recruits SHREC during the S phase. We also showed that the HBS domain, which is involved in the repression of early replication origins (36), is essential for SHREC localization (Fig. 5D). Furthermore, we showed that Mrc1 physically interacts with Clr3 via Mrc1’s HBS containing C-terminal region (Fig. 6A-6C). These data suggest that HBS acts as a physical interaction surface between Mrc1 and SHREC. Another possibility is that HBS-mediated control of replication timing may play an important role in the recruitment of SHREC, although replication origins located within subtelomeres fire at the late S phase (48, 49). In our opinion, in addition to temporally and spatially, the timely recruitment of SHREC during the S phase by Mrc1 may be important in maintaining heterochromatin structure.

### Coordination of the DNA replication and heterochromatin maintenance

The onset of DNA replication is regulated by histone acetylation. In human cells, the MYST family histone acetyltransferase, HBO1, has been reported to be associated with DNA replication initiation factors (50, 51), and acetylation of histone H3K14 facilitates efficient activation of DNA replication (52). In budding yeast, the Rpd3p histone deacetylase delays the firing of the replication origin, whereas the Gcn5p histone acetyltransferase accelerates the firing of the replication start site (53). ChIP analysis showed that the level of H3K14ac at the subtelomere was elevated in the early/mid S phase (Fig. 5E). Therefore, H3K14ac is likely important for the initiation of DNA replication in *S. pombe*. In the heterochromatin region, elevated H3K14ac levels must rapidly decrease to repress transcription. Mrc1 localized on chromatin during the S phase when the level of H3K14ac began to decrease and that of H3K9me2 began to increase (Fig. 5E). This suggests that Mrc1 may function to rapidly reduce H3K14ac levels at the end of the S phase. In budding yeast, Mrc1 interacts with DNA polymerase ε, Pol2 (54). In the fission yeast, DNA polymerase ε, Cdc20 forms a complex with Rik1 and Dos2, components of CLRC, and play important roles in the maintenance of H3K9me in heterochromatin (55). In addition, Dpb3-Dpb4, which forms a complex with Cdc20, promotes H3K9 hypoacetylation, together with two HDACs, Clr6 and Sir2 (41). However, the relationship between SHREC, which promotes the deacetylation of histone H3K14, and DNA replication factors remains unclear. We found that Mrc1, which functions in replication forks, promotes the maintenance of the hypoacetylated state of H3K14 through the recruitment of SHREC (Fig. 5A, 5D). These findings suggest that factors associated with DNA replication forks have multi-layered functions in the inheritance of repressive histone modifications (Fig. 8). In mammals, <u>P</u>roliferating <u>C</u>ell <u>N</u>uclear <u>A</u>ntigen (PCNA) at the replication fork recruits G9a, an H3K9 methyltransferase, and HDAC2, a histone deacetylase that promotes epigenetic inheritance of repressive histone modifications (56). Our findings shed light on the evolutionarily conserved mechanisms of epigenetic control mediated by DNA replication fork components in eukaryotes.

**Fig. 8.**
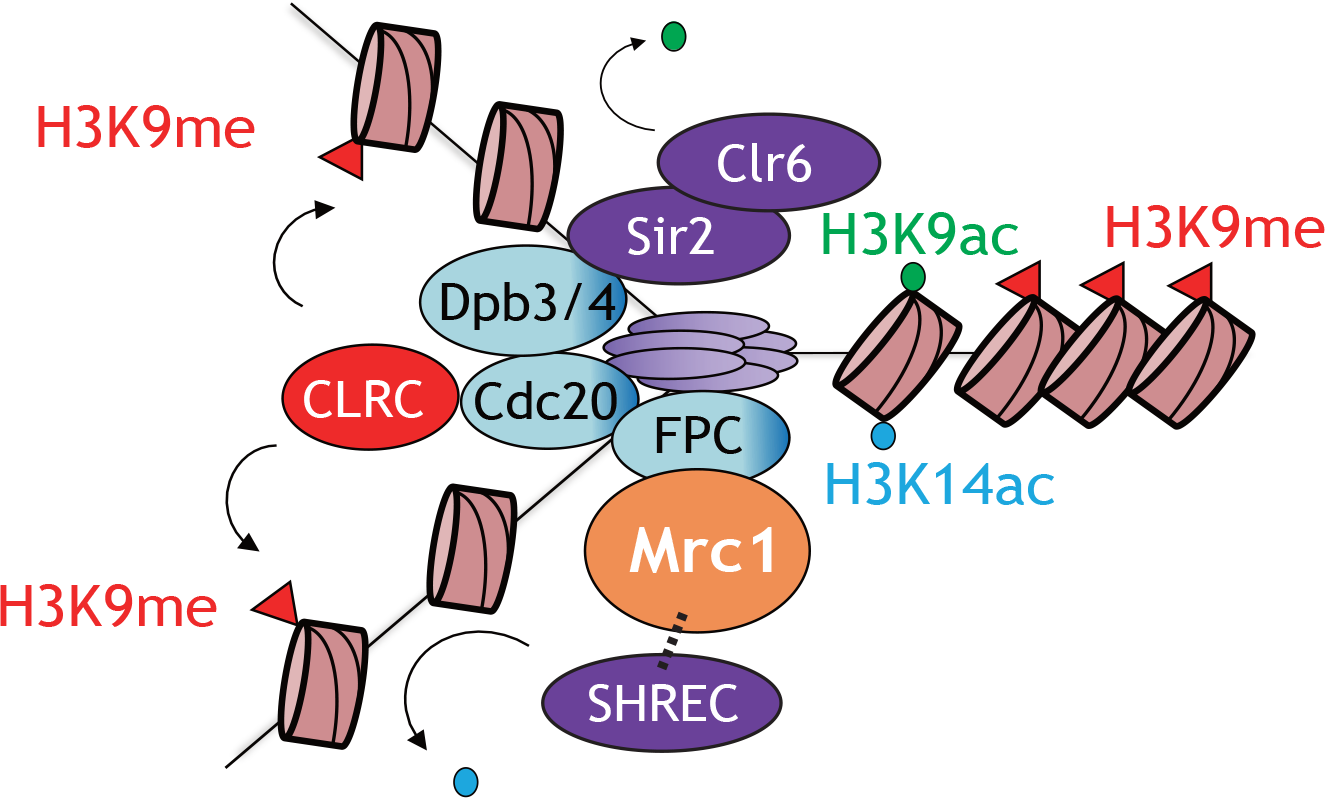
Model of this study: Mrc1 maintains heterochromatin through the recruitment of SHREC, a H3K14 deacetyltransferase complex At the onset of DNA replication, H3K9 is demethylated, and H3K9 and K14 are transiently acetylated in heterochromatin. During DNA replication, Cdc20 accessory proteins, Dpb3-Dpb4 recruit Sir2 and Clr6, resulting in the deacetylation of H3K9. Simultaneously, Mrc1 recruits SHREC to the replication fork to promote H3K14 deacetylation. These histone deacetylation reactions allow CLRC to methylate H3K9 effectively.

## Materials and Methods

### *S. pombe* strains, media, and growth conditions

Except for annotation, the cells were cultured in YES medium at 30 °C. 5-FOA was added to YES at a final concentration of 0.1 % (w/v). All deletion mutants or tagged strains were generated using PCR-based techniques or crossings (57, 58). Gene deletion and epitope tagging was confirmed using PCR and western blotting. The strains used in this study are listed in supplemental table (Table. S1).

### Spot assay

Overnight cultures in YES medium were harvested, and a 1×10^8^ cells/mL suspension was prepared with dH2O. Then, a five-fold dilution series of cell suspensions was prepared and spotted on YES, YE, or YES + 0.1 % 5-FOA plates. Images were taken after three days of incubation.

### Systematic screening with *S. pombe* haploid deletion library

For the primary screening, a tester strain harboring *subtel*::*ade6*^+^ (Bio5668) was crossed with *S. pombe* haploid deletion library mutants (ver 3.0, Bioneer, Daejeon, South Korea) and an additional gene deletion set that was previously generated by the K. Gould Lab on SPAS for 3 d, followed by treatment with 30 % EtOH to kill vegetative cells (28, 29). Spores were spotted onto YE plates containing G418, hygromycin B, and clonNAT, and were incubated for 3–4 days at 30 °C to assess colony color. For secondary screening, a tester strain harboring *subtel*::*ura4*^+^ (KH6380) was crossed with candidates that passed the primary screening. After sporulation, at least two G418- and clonNAT resistant progenies were spotted on YES plates containing 0.1 % 5-FOA for testing their sensitivity to 5-FOA. 5-FOA sensitive strains were identified as subtelomeric heterochromatin mutants (Table. 1).

### RNA preparation and RT-PCR analysis

Total RNA was extracted from 5×10^7^ exponentially growing cells using the hot phenol method as described previously (59). Reverse transcription and quantitative PCR were performed in one step using the Luna Universal One-Step RT-qPCR Kit (New England BioLabs, Ipswich, MA, USA). The primers used are listed in supplemental table (Table. S2).

### ChIP analysis

1–2.5×10^8^ exponentially growing cells (cultured in YES medium at 30 ℃) were fixed with 1 % formaldehyde (nacalai tesque, Kyoto, Japan) for 30 min at 25 ℃ and quenched by the addition of 150 mM glycine. Cells were harvested and washed three times with Buf.1 (50 mM HEPES, 140 mM NaCl, 1 mM EDTA, 1 % TritonX-100, 0.1 % sodium deoxycholate, pH 7.5). Cells were resuspended in Buf1*(10 ml Buf.1 + protease inhibitor cocktail (Roche, Basel, Switzerland) and lysed with MULTI BEADS SHOCKER (YASUI KIKAI, Osaka, Japan). To shear chromatin, the total lysate was sonicated for 7.5 min (30 s ON, 30 s OFF, 15 cycles) at level H (250 W) using a Bioruptor II (Sonicbio, Kanagawa, Japan). The soluble fraction obtained by centrifugation at 15,000 rpm for 15 min was reacted with sheep anti-mouse or anti-rabbit primary antibody-conjugated Dynabeads M280 (Thermo Fisher Scientific, Waltham, MA, USA) for 4 h or overnight. Anti-H3K9me2 (5.1.1, mAbProtein, Shimane, Japan), anti-H3K14ac (EP964Y, abcam, Cambridge, UK), anti-Swi6 (Gift from Dr. S. Takahata), anti-Chp2 (Gift from Dr. J. Nakayama), anti-FLAG (M2, Sigma-Aldrich, St. Louis, MO, USA), and anti-myc (9B11, Cell signaling Technology, Danvers, MA, USA) were used as primary antibodies. Beads were washed twice each with Buf.1, Buf.1’ (50 mM HEPES, 500 mM NaCl, 1 mM EDTA, 1 % TritonX-100, 0.1 % sodium deoxycolate, pH 7.5), Buf.2 (10 mM Tris, 250 mM LiCl, 0.5 % NP-40, 0.5 % sodium deoxycholate, pH 8.0), and TE. Washed beads were suspended in reversal buffer (20 mM Tris, 1 mM EDTA, 0.8 % SDS, pH 8.0) and incubated for 15 min at 95 ℃. DNA was purified using the FastGene Gel/PCR Extraction Kit (NIPPON Genetics, Tokyo, Japan). Purified DNA was quantified using GeneAce SYBR qPCR Mix α (NIPPON GENE, Tokyo, Japan) combined with a LightCycler 96 Real-Time PCR system (Roche, Basel, Switzerland). The primers used are listed in supplemental table (Table. S2).

### Cell cycle synchronization and ChIP analysis by *nda3-KM311*

Cells harboring the cold-sensitive mutation, *nda3-KM311*, were grown to 2×10^6^ cells/mL at 28 °C. The cells were arrested at prophase by cooling the culture to 20 °C and incubating for 4–5 h. The culture was then quickly transferred to a 28 °C incubator to restart the cell cycle. Every 10 min, 6 × 10^7^ cells were collected and fixed in 1 % formaldehyde at 25 °C for 20 min. Subsequent experimental procedures were performed as described for the ChIP analysis.

### Yeast two-hybrid assay

pGBT9-mrc1 (pKT3) was constructed by cloning the full-length *mrc1* coding region into the EcoRI-SmaI double-digested pGBT9 vector. pGBKT7-mrc1-N (pKT67), pGBKT7-mrc1-M (pKT68), or pGBKT7-mrc1-C (pKT69) was constructed by cloning the N-terminal region (1-503 a.a.), the middle region (248-756 a.a.), or the C-terminal region (500-1019 a.a.) of Mrc1, respectively, into the EcoRI-PstI double-digested pGBKT7 vector. pGADT7-clr1 (pKT3216), pGADT7-clr2 (pKT3217), pGADT7-clr3 (pKT3218), and pGADT7-mit1 (pKT3219) were constructed by cloning the full-length coding region into the pGADT7 vector using the In-Fusion HD Cloning Kit (Clontech Laboratories, Mountain View, CA, USA). The Matchmaker two-hybrid system 3 (Clontech Laboratories, Mountain View, CA, USA) was used for the yeast two-hybrid assay, according to the manufacturer’s instructions. The interaction was scored by monitoring growth on SC-LEU-TRP (SC-LW), SC-LEU-TRP-HIS (SC-LWH) + 20 mM 3-Amino-1,2,4-triazole (3-AT), or SC-LEU-TRP-HIS (SC-LWH) + 100 mM 3-AT plates. The plasmids used are listed in supplemental table (Table. S3).

## Acknowledgement

We thank Dr. H. Masai, Dr. R. Allshire, and Dr. SI. Grewal, and Dr. Y. Murakami for providing the strains, and Dr. J. Nakayama, Dr. T. Urano, and Dr. S. Takahata for providing antibodies. We would also like to thank Dr. H. Masai and Dr. G. Thon for discussion and sharing unpublished data, and Dr. J. Nakayama for critically reading the manuscript. We would like to thank all members of the Tanaka Lab for their support. This study was supported by the Foundation of Kinoshita Memorial Enterprise (K. K.) and JSPS KAKENHI (grant number JP20H02952 to K. T.).

## Author Approvals

All authors have seen and approved the manuscript. This manuscript hasn’t been accepted or published elsewhere.

## Competing Interests

The authors declare no competing interest.

## Notes

### Competing Interest Statement

The authors have declared no competing interest.

## References

1. Jenuwein T, Allis CD. Translating the histone code. Science. 2001;293(5532):1074–80.

2. Turner BM. Cellular memory and the histone code. Cell. 2002;111(3):285–91.

3. Padeken J, Methot SP, Gasser SM. Establishment of H3K9- methylated heterochromatin and its functions in tissue differentiation and maintenance. Nat Rev Mol Cell Biol. 2022;23(9):623–40.

4. Volpe TA, Kidner C, Hall IM, Teng G, Grewal SI, Martienssen RA. Regulation of heterochromatic silencing and histone H3 lysine-9 methylation by RNAi. Science. 2002;297(5588):1833-7.

5. Allshire RC, Ekwall K. Epigenetic Regulation of Chromatin States in Schizosaccharomyces pombe. Cold Spring Harb Perspect Biol. 2015;7(7):a018770.

6. Goto DB, Nakayama J. RNA and epigenetic silencing: insight from fission yeast. Dev Growth Differ. 2012;54(1):129–41.

7. Verdel A, Jia S, Gerber S, Sugiyama T, Gygi S, Grewal SI, et al. RNAi-mediated targeting of heterochromatin by the RITS complex. Science. 2004;303(5658):672-6.

8. Nakayam J, Rice JC, Strahl BD, Allis CD, Grewal SIS. Role of histone H3 lysine 9 methylation in epigenetic control of heterochromatin assembly. Science. 2001;292(5514):110-3.

9. Zhang K, Mosch K, Fischle W, Grewal SI. Roles of the Clr4 methyltransferase complex in nucleation, spreading and maintenance of heterochromatin. Nat Struct Mol Biol. 2008;15(4):381–8.

10. Sugiyama T, Cam HP, Sugiyama R, Noma K, Zofall M, Kobayashi R, et al. SHREC, an effector complex for heterochromatic transcriptional silencing. Cell. 2007;128(3):491–504.

11. Motamedi MR, Hong EJ, Li X, Gerber S, Denison C, Gygi S, et al. HP1 proteins form distinct complexes and mediate heterochromatic gene silencing by nonoverlapping mechanisms. Mol Cell. 2008;32(6):778–90.

12. Fischer T, Cui BW, Dhakshnamoorthy J, Zhou M, Rubin C, Zofall M, et al. Diverse roles of HP1 proteins in heterochromatin assembly and functions in fission yeast. Proceedings of the National Academy of Sciences of the United States of America. 2009;106(22):8998–9003.

13. Sadaie M, Kawaguchi R, Ohtani Y, Arisaka F, Tanaka K, Shirahige K, et al. Balance between Distinct HP1 Family Proteins Controls Heterochromatin Assembly in Fission Yeast. Molecular and Cellular Biology. 2008;28(23):6973–88.

14. Hall IM, Shankaranarayana GD, Noma K, Ayoub N, Cohen A, Grewal SI. Establishment and maintenance of a heterochromatin domain. Science. 2002;297(5590):2232-7.

15. Audergon PN, Catania S, Kagansky A, Tong P, Shukla M, Pidoux AL, et al. Epigenetics. Restricted epigenetic inheritance of H3K9 methylation. Science. 2015;348(6230):132-5.

16. Ragunathan K, Jih G, Moazed D. Epigenetics. Epigenetic inheritance uncoupled from sequence-specific recruitment. Science. 2015;348(6230):1258699.

17. Grewal SIS, Klar AJS. Chromosomal inheritance of epigenetic states in fission yeast during mitosis and meiosis. Cell. 1996;86(1):95–101.

18. Folco HD, McCue A, Balachandran V, Grewal SIS. Cohesin Impedes Heterochromatin Assembly in Fission Yeast Cells Lacking Pds5. Genetics. 2019;213(1):127–41.

19. Shipkovenska G, Durango A, Kalocsay M, Gygi SP, Moazed D. A conserved RNA degradation complex required for spreading and epigenetic inheritance of heterochromatin. Elife. 2020;9.

20. Iglesias N, Paulo JA, Tatarakis A, Wang X, Edwards AL, Bhanu NV, et al. Native Chromatin Proteomics Reveals a Role for Specific Nucleoporins in Heterochromatin Organization and Maintenance. Mol Cell. 2020;77(1):51–66.e8.

21. Taneja N, Zofall M, Balachandran V, Thillainadesan G, Sugiyama T, Wheeler D, et al. SNF2 Family Protein Fft3 Suppresses Nucleosome Turnover to Promote Epigenetic Inheritance and Proper Replication. Mol Cell. 2017;66(1):50–62.e6.

22. Holla S, Dhakshnamoorthy J, Folco HD, Balachandran V, Xiao H, Sun LL, et al. Positioning Heterochromatin at the Nuclear Periphery Suppresses Histone Turnover to Promote Epigenetic Inheritance. Cell. 2020;180(1):150–64.e15.

23. Kanoh J, Sadaie M, Urano T, Ishikawa F. Telomere binding protein Taz1 establishes Swi6 heterochromatin independently of RNAi at telomeres. Curr Biol. 2005;15(20):1808–19.

24. Hansen KR, Ibarra PT, Thon G. Evolutionary-conserved telomere- linked helicase genes of fission yeast are repressed by silencing factors, RNAi components and the telomere-binding protein Taz1. Nucleic Acids Res. 2006;34(1):78–88.

25. Barrales RR, Forn M, Georgescu PR, Sarkadi Z, Braun S. Control of heterochromatin localization and silencing by the nuclear membrane protein Lem2. Genes Dev. 2016;30(2):133–48.

26. Ekwall K, Olsson T, Turner BM, Cranston G, Allshire RC. Transient inhibition of histone deacetylation alters the structural and functional imprint at fission yeast centromeres. Cell. 1997;91(7):1021–32.

27. Cam HP, Sugiyama T, Chen ES, Chen X, FitzGerald PC, Grewal SI. Comprehensive analysis of heterochromatin- and RNAi-mediated epigenetic control of the fission yeast genome. Nat Genet. 2005;37(8):809–19.

28. Chen JS, Beckley JR, McDonald NA, Ren L, Mangione M, Jang SJ, et al. Identification of new players in cell division, DNA damage response, and morphogenesis through construction of Schizosaccharomyces pombe deletion strains. G3 (Bethesda). 2014;5(3):361-70.

29. Chen JS, Beckley JR, Ren L, Feoktistova A, Jensen MA, Rhind N, et al. Discovery of genes involved in mitosis, cell division, cell wall integrity and chromosome segregation through construction of Schizosaccharomyces pombe deletion strains. Yeast. 2016;33(9):507–17.

30. Jahn LJ, Mason B, Brøgger P, Toteva T, Nielsen DK, Thon G. Dependency of Heterochromatin Domains on Replication Factors. G3 (Bethesda). 2018;8(2):477-89.

31. Tanaka K, Russell P. Mrc1 channels the DNA replication arrest signal to checkpoint kinase Cds1. Nature Cell Biology. 2001;3(11):966–72.

32. Masai H, Yang CC, Matsumoto S. Mrc1/Claspin: a new role for regulation of origin firing. Current Genetics. 2017;63(5):813–8.

33. Zhao H, Russell P. DNA binding domain in the replication checkpoint protein Mrc1 of Schizosaccharomyces pombe. Journal of Biological Chemistry. 2004;279(51):53023–7.

34. Tanaka T, Yokoyama M, Matsumoto S, Fukatsu R, You ZY, Masai H. Fission Yeast Swi1-Swi3 Complex Facilitates DNA Binding of Mrc1. Journal of Biological Chemistry. 2010;285(51):39609–22.

35. Zhao H, Tanaka K, Nogochi E, Nogochi C, Russell P. Replication checkpoint protein Mrc1 is regulated by Rad3 and Tel1 in fission yeast. Molecular and Cellular Biology. 2003;23(22):8395–403.

36. Matsumoto S, Kanoh Y, Shimmoto M, Hayano M, Ueda K, Fukatsu R, et al. Checkpoint-Independent Regulation of Origin Firing by Mrc1 through Interaction with Hsk1 Kinase. Molecular and Cellular Biology. 2017;37(7).

37. Grewal SI, Klar AJ. A recombinationally repressed region between mat2 and mat3 loci shares homology to centromeric repeats and regulates directionality of mating-type switching in fission yeast. Genetics. 1997;146(4):1221–38.

38. Toda T, Umesono K, Hirata A, Yanagida M. Cold-sensitive nuclear division arrest mutants of the fission yeast Schizosaccharomyces pombe. J Mol Biol. 1983;168(2):251–70.

39. Hiraoka Y, Toda T, Yanagida M. The NDA3 gene of fission yeast encodes beta-tubulin: a cold-sensitive nda3 mutation reversibly blocks spindle formation and chromosome movement in mitosis. Cell. 1984;39(2 Pt 1):349-58.

40. Chen ES, Zhang K, Nicolas E, Cam HP, Zofall M, Grewal SI. Cell cycle control of centromeric repeat transcription and heterochromatin assembly. Nature. 2008;451(7179):734-7.

41. He H, Li Y, Dong Q, Chang AY, Gao F, Chi Z, et al. Coordinated regulation of heterochromatin inheritance by Dpb3-Dpb4 complex. Proc Natl Acad Sci U S A. 2017;114(47):12524–9.

42. Kato H, Okazaki K, Iida T, Nakayama J, Murakami Y, Urano T. Spt6 prevents transcription-coupled loss of posttranslationally modified histone H3. Scientific Reports. 2013;3.

43. Chikashige Y, Yamane M, Okamasa K, Tsutsumi C, Kojidani T, Sato M, et al. Membrane proteins Bqt3 and -4 anchor telomeres to the nuclear envelope to ensure chromosomal bouquet formation. J Cell Biol. 2009;187(3):413–27.

44. Funabiki H, Hagan I, Uzawa S, Yanagida M. Cell cycle-dependent specific positioning and clustering of centromeres and telomeres in fission yeast. J Cell Biol. 1993;121(5):961–76.

45. Mizuguchi T, Fudenberg G, Mehta S, Belton JM, Taneja N, Folco HD, et al. Cohesin-dependent globules and heterochromatin shape 3D genome architecture in S. pombe. Nature. 2014;516(7531):432-5.

46. Mizuguchi T, Barrowman J, Grewal SI. Chromosome domain architecture and dynamic organization of the fission yeast genome. FEBS Lett. 2015;589(20 Pt A):2975-86.

47. Shimada A, Dohke K, Sadaie M, Shinmyozu K, Nakayama J, Urano T, et al. Phosphorylation of Swi6/HP1 regulates transcriptional gene silencing at heterochromatin. Genes Dev. 2009;23(1):18–23.

48. Hayashi M, Katou Y, Itoh T, Tazumi A, Tazumi M, Yamada Y, et al. Genome-wide localization of pre-RC sites and identification of replication origins in fission yeast. EMBO J. 2007;26(5):1327–39.

49. Hayashi MT, Takahashi TS, Nakagawa T, Nakayama J, Masukata H. The heterochromatin protein Swi6/HP1 activates replication origins at the pericentromeric region and silent mating-type locus. Nat Cell Biol. 2009;11(3):357–62.

50. Iizuka M, Stillman B. Histone acetyltransferase HBO1 interacts with the ORC1 subunit of the human initiator protein. J Biol Chem. 1999;274(33):23027–34.

51. Burke TW, Cook JG, Asano M, Nevins JR. Replication factors MCM2 and ORC1 interact with the histone acetyltransferase HBO1. J Biol Chem. 2001;276(18):15397–408.

52. Feng Y, Vlassis A, Roques C, Lalonde ME, González-Aguilera C, Lambert JP, et al. BRPF3-HBO1 regulates replication origin activation and histone H3K14 acetylation. EMBO J. 2016;35(2):176–92.

53. Vogelauer M, Rubbi L, Lucas I, Brewer BJ, Grunstein M. Histone acetylation regulates the time of replication origin firing. Mol Cell. 2002;10(5):1223–33.

54. Lou H, Komata M, Katou Y, Guan Z, Reis CC, Budd M, et al. Mrc1 and DNA polymerase epsilon function together in linking DNA replication and the S phase checkpoint. Mol Cell. 2008;32(1):106–17.

55. Li F, Martienssen R, Cande WZ. Coordination of DNA replication and histone modification by the Rik1-Dos2 complex. Nature. 2011;475(7355):244-8.

56. Groth A, Rocha W, Verreault A, Almouzni G. Chromatin challenges during DNA replication and repair. Cell. 2007;128(4):721–33.

57. Krawchuk MD, Wahls WP. High-efficiency gene targeting in Schizosaccharomyces pombe using a modular, PCR-based approach with long tracts of flanking homology. Yeast. 1999;15(13):1419–27.

58. Bähler J, Wu JQ, Longtine MS, Shah NG, McKenzie A, Steever AB, et al. Heterologous modules for efficient and versatile PCR-based gene targeting in Schizosaccharomyces pombe. Yeast. 1998;14(10):943–51.

59. Kawakami K, Hayashi A, Nakayama J, Murakami Y. A novel RNAi protein, Dsh1, assembles RNAi machinery on chromatin to amplify heterochromatic siRNA. Genes Dev. 2012;26(16):1811–24.

