## Supplemental Fig_1 for "Mrc1^Claspin^ is essential for heterochromatin maintenance in *Schizosaccharomyces pombe*"

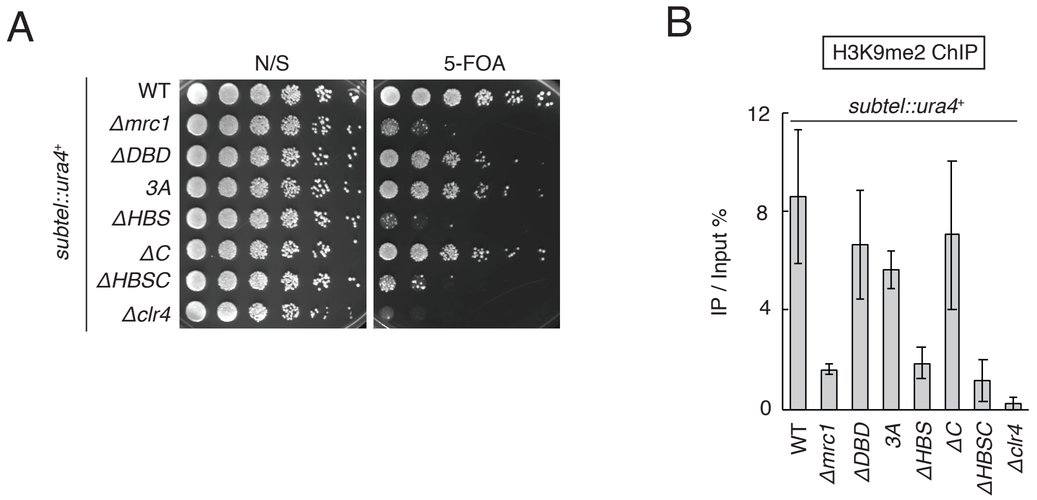


**Fig. S1 HBS is required for Mrc1's silencing function**

***(A)*** Five-fold serial dilutions of indicated genotype cells harboring *subtel*::*ura4*$^{+}$ were spotted onto YES and YES + 0.1 % 5-FOA plates and incubated for 3 d at 30 ℃. ***(B)*** H3K9me2 ChIP at *subtel*::*ura4*$^{+}$ in strains used in *(A)*. The vertical axis shows the IP %. Error bars show the standard deviation from three independent cultures.
