## Supplementary material for "Mrc1^Claspin^ is essential for heterochromatin maintenance in *Schizosaccharomyces pombe*": Table_S1

### Table S1: Strain list

| Strain | Genotype | Figure | Origin |
|---|---|---|---|
| **PR109** | $h^{-}$ leu1-32 ura4-D18 | 1B | Dr. P. Russell |
| **FY2002** | $h^{+}$ leu1-32 ura4-DS/E ade6-DN/N imr1L::$ura4^{+}$ otr1R::$ade6^{+}$ | 1B, 4A | Dr. R. Allshire |
| **Bio5666** | $h^{-}$ leu1-32 ura4-D18 Δade6::hphMX6 SPAC212.12 5'UTR::$ade6^{+}$ (red and white mixture) | 1B-1F, 2A | This study |
| **Bio5667** | $h^{-}$ leu1-32 ura4-D18 Δade6::hphMX6 SPAC212.12 5'UTR::$ade6^{+}$ (red isolate) | 1G | This study |
| **Bio5668** | $h^{-}$ leu1-32 ura4-D18 Δade6::hphMX6 SPAC212.12 5'UTR::$ade6^{+}$ nhe1-GFP-natMX6 (red isolate) | 2B | This study |
| **Bio5665** | $h^{+}$ leu1-32 ura4-D18 Δade6::hphMX6 SPAC212.12 5'UTR::$ade6^{+}$ Δclr4::kanMX6 | 1G | This study |
| **Bio9876** | $h^{?}$ leu1-32 ura4-D18 Δade6::hphMX6 SPAC212.12 5'UTR::$ade6^{+}$ nhe1-GFP-natMX6 Δswi6::kanMX4 | 1G | This study |
| **Bio9878** | $h^{?}$ leu1-32 ura4-D18 Δade6::hphMX6 SPAC212.12 5'UTR::$ade6^{+}$ nhe1-GFP-natMX6 Δchp2::kanMX4 | 1G | This study |
| **Bio9877** | $h^{?}$ leu1-32 ura4-D18 Δade6::hphMX6 SPAC212.12 5'UTR::$ade6^{+}$ nhe1-GFP-natMX6 Δclr3::kanMX4 | 1G | This study |
| **Bio7622** | $h^{-}$ leu1-32 ura4-D18 Δade6::hphMX6 SPAC212.12 5'UTR::$ade6^{+}$ nhe1-GFP-natMX6 Δago1::bleMX6 | 1G | This study |
| **KK5477** | $h^{+}$ leu1-32 ura4-DS/E ade6-DN/N imr1L::$ura4^{+}$ otr1R::$ade6^{+}$ Δclr4::kanMX6 | 1G, 4A | Dr. Y. Murakami |
| **Bio8833** | $h^{+}$ leu1-32 ura4-DS/E ade6-DN/N imr1L::$ura4^{+}$ otr1R::$ade6^{+}$ Δswi6::hphMX6 | 1G | Dr. Y. Murakami |
| **Bio8834** | $h^{+}$ leu1-32 ura4-DS/E ade6-DN/N imr1L::$ura4^{+}$ otr1R::$ade6^{+}$ Δchp2::kanMX6 | 1G | Dr. Y. Murakami |
| **Bio8832** | $h^{+}$ leu1-32 ura4-DS/E ade6-DN/N imr1L::$ura4^{+}$ otr1R::$ade6^{+}$ Δclr3::hphMX6 | 1G | Dr. Y. Murakami |
| **Bio8739** | $h^{+}$ leu1-32 ura4-DS/E ade6-DN/N imr1L::$ura4^{+}$ otr1R::$ade6^{+}$ Δago1::hphMX6 | 1G, 4A | This study |
| **KH6380** | $h^{-}$ leu1-32 ura4-D18 SPAC212.07::$ura4^{+}$ nhe1-GFP-natMX6 | 2C | This study |
| **Bio7632** | $h^{?}$ leu1-32 ura4-D18 Δade6::hphMX6 SPAC212.12 5'UTR::$ade6^{+}$ nhe1-GFP-natMX6 Δmrc1::kanMX6 | 3B-3D | This study |
| **Bio7633** | $h^{?}$ leu1-32 ura4-D18 Δade6::hphMX6 SPAC212.12 5'UTR::$ade6^{+}$ nhe1-GFP-natMX6 mrc1Δ782-879-3FLAG::kanMX6 (ΔHBS) | 3B-3D | This study |
| **Bio7634** | $h^{?}$ leu1-32 ura4-D18 Δade6::hphMX6 SPAC212.12 5'UTR::$ade6^{+}$ nhe1-GFP-natMX6 mrc1Δ880-1019-5FLAG::kanMX6 (ΔC) | 3B-3D | This study |
| **Bio7635** | $h^{?}$ leu1-32 ura4-D18 Δade6::hphMX6 SPAC212.12 5'UTR::$ade6^{+}$ nhe1-GFP-natMX6 mrc1Δ782-1019-5FLAG::kanMX6 (ΔHBSC) | 3B-3D | This study |
| **Bio7636** | $h^{?}$ leu1-32 ura4-D18 Δade6::hphMX6 SPAC212.12 5'UTR::$ade6^{+}$ nhe1-GFP-natMX6 Δcds1::kanMX6 | 3B-3D | This study |
| **Bio7637** | $h^{?}$ leu1-32 ura4-D18 Δade6::hphMX6 SPAC212.12 5'UTR::$ade6^{+}$ nhe1-GFP-natMX6 mrc1Δ160-284-13myc::kanMX6 (ΔDBD) | 3B-3D | This study |
| **Bio7638** | $h^{?}$ leu1-32 ura4-D18 Δade6::hphMX6 SPAC212.12 5'UTR::$ade6^{+}$ nhe1-GFP-natMX6 mrc1-S604AT645AT653A-5FLAG::kanMX6 (3A) | 3B-3D | This study |
| **NH5754** | $h^{+}$ leu1-32 ura4-D18 SPAC212.07::$ura4^{+}$ | S1, 5A, 5B, 5C, 7A, 7B | This study |
| **NH5755** | $h^{+}$ leu1-32 ura4-D18 SPAC212.07::$ura4^{+}$ Δmrc1::kanMX6 | S1, 4B, 5A, 5B, 5C, 7A, 7B | This study |

| | | | |
|---|---|---|---|
| **Bio6060** | *h*$^{?}$ *leu1-32 ura4-D18 SPAC212.07::ura4*$^{+}$ *nhe1-GFP-natMX6 mrc1Δ160-284-13myc::kanMX6* (*ΔDBD*) | S1 | This study |
| **YU6337** | *h*$^{?}$ *leu1-32 ura4-D18 SPAC212.07::ura4*$^{+}$ *mrc1-S604AT645AT653A-5FLAG::kanMX6* (*3A*) | S1 | This study |
| **Bio5985** | *h*$^{?}$ *leu1-32 ura4-D18 SPAC212.07::ura4*$^{+}$ *mrc1Δ782-879-3FLAG::kanMX6* (*ΔHBS*) | S1, 4B, 5A, 5B, 5C, 7A, 7B | This study |
| **YU6608** | *h*$^{+}$ *leu1-32 ura4-D18 SPAC212.07::ura4*$^{+}$ *mrc1Δ880-1019-5FLAG::kanMX6* (*ΔC*) | S1 | This study |
| **YU6610** | *h*$^{+}$ *leu1-32 ura4-D18 SPAC212.07::ura4*$^{+}$ *mrc1Δ782-1019-5FLAG::kanMX6* (*ΔHBSC*) | S1 | This study |
| **NH5761** | *h*$^{+}$ *leu1-32 ura4-D18 SPAC212.07::ura4*$^{+}$ *Δclr4::kanMX6* | S1, 2C, 5A, 5B, 5C | This study |
| **NH5776** | *h*$^{+}$ *leu1-32 ura4-DS/E ade6-DN/N imr1L::ura4*$^{+}$ *otr1R::ade6*$^{+}$*Δmrc1::kanMX6* | 4A | This study |
| **Bio7361** | *h*$^{+}$ *leu1-32 ura4-D18 SPAC212.07::ura4*$^{+}$ *Δago1::hphMX6* | 4B | This study |
| **Bio7362** | *h*$^{?}$ *leu1-32 ura4-D18 SPAC212.07::ura4*$^{+}$ *Δago1::hphMX6 mrc1Δ782-879-3FLAG::kanMX6* (*ΔHBS*) | 4B | This study |
| **Bio7363** | *h*$^{+}$ *leu1-32 ura4-D18 SPAC212.07::ura4*$^{+}$ *Δago1::hphMX6 Δmrc1::kanMX6* | 4B | This study |
| **KT693** | *h*$^{90}$ *leu1-32 ura4-DS/E ade6-m210 his2 Kint2::ura4*$^{+}$ *Δclr4::kanMX6* | 4C, 4D | Lab stock |
| **KT694** | *h*$^{90}$ *leu1-32 ura4-DS/E ade6-m210 his2 Kint2::ura4*$^{+}$ | 4C, 4D | Dr. SI. Grewal |
| **YU7299** | *h*$^{90}$ *leu1-32 ura4 ade6? his2 Kint2::ura4*$^{+}$ *Δmrc1::kanMX6* | 4C, 4D | This study |
| **YU7301** | *h*$^{90}$ *leu1-32 ura4 ade6? his2 Kint2::ura4*$^{+}$ *mrc1-S604AT645AT653A-5FLAG::kanMX6* (*3A*) | 4C, 4D | This study |
| **YU7303** | *h*$^{90}$ *leu1-32 ura4 ade6? his2 Kint2::ura4*$^{+}$ *mrc1Δ160-284-5FLAG::kanMX6* (*ΔDBD*) | 4C, 4D | This study |
| **YU7305** | *h*$^{90}$ *leu1-32 ura4 ade6? his2 Kint2::ura4*$^{+}$ *mrc1Δ782-879-3FLAG::kanMX6* (*ΔHBS*) | 4C, 4D | This study |
| **YU7383** | *h*$^{90}$ *leu1-32 ura4-DS/E ade6-m210 his2 Kint2::ura4*$^{+}$ *mrc1Δ782-1019-5FLAG::kanMX6* (*ΔHBSC*) | 4C, 4D | This study |
| **YU7386** | *h*$^{90}$ *leu1-32 ura4-DS/E ade6-m210 his2 Kint2::ura4*$^{+}$ *mrc1Δ880-1019-5FLAG::kanMX6* (*ΔC*) | 4C, 4D | This study |
| **NH6076** | *h*$^{-}$ *leu1-32 ura4-D18 ade6-m210 his2 ΔK::ura4*$^{+}$ | 4C, 4D, 7C, 7D | Dr. SI. Grewal |
| **NH6083** | *h*$^{-}$ *leu1-32 ura4-D18 ade6-m210 his2 ΔK::ura4*$^{+}$ *Δclr4::kanMX6* | 4C, 4D | This study |
| **YU6673** | *h*$^{-}$ *leu1-32 ura4-D18 ade6? Δhis2::natMX6 ΔK::ura4*$^{+}$ *mrc1-S604AT645AT653A-5FLAG::kanMX6* (*3A*) | 4C, 4D | This study |
| **YU6683** | *h*$^{-}$ *leu1-32 ura4-D18 ade6? Δhis2::natMX6 ΔK::ura4*$^{+}$ *mrc1Δ160-284-13myc::kanMX6* (*ΔDBD*) | 4C, 4D | This study |
| **KT6927** | *h*$^{-}$ *leu1-32 ura4-D18 ade6-m210 his2 ΔK::ura4*$^{+}$ *Δmrc1::kanMX6* | 4C, 4D, 7C, 7D | This study |
| **YU7308** | *h*$^{-}$ *leu1-32 ura4-D18 ade6-m210 his2 ΔK::ura4*$^{+}$ *mrc1Δ782-879-3FLAG::kanMX6* (*ΔHBS*) | 4C, 4D | This study |
| **YU7310** | *h*$^{-}$ *leu1-32 ura4-D18 ade6-m210 his2 ΔK::ura4*$^{+}$ *mrc1Δ782-1019-5FLAG::kanMX6* (*ΔHBSC*) | 4C, 4D, 7C, 7D | This study |
| **YU7312** | *h*$^{-}$ *leu1-32 ura4-D18 ade6-m210 his2 ΔK::ura4*$^{+}$ *mrc1Δ880-1019-5FLAG::kanMX6* (*ΔC*) | 4C, 4D | This study |
| **KT260** | *h*$^{-}$ *leu1-32 ura4-D18 nda3-KM311* | 5D, 5E | Lab stock |
| **KT1641** | *h*$^{-}$ *leu1-32 ura4-D18 nda3-KM311 mrc1-5FLAG::kanMX6* | 5E | This study |
| **Bio7321** | *h*$^{-}$ *leu1-32 ura4-D18 nda3-KM311 mit1-13myc::hphMX6* | 5D | This study |

| | | | |
|---|---|---|---|
| **Bio7323** | $h^{?}$ *leu1-32 ura4-D18 nda3-KM311 mit1-13myc*::*natMX6* Δ*mrc1*::*kanMX6* | 5D | This study |
| **Bio7397** | $h^{?}$ *leu1-32 ura4-D18 nda3-KM311 mit1-13myc*::*hphMX6 mrc1*Δ*782-879-3FLAG*::*kanMX6* (Δ*HBS*) | 5D | This study |
| **Bio7061** | $h^{+}$ *leu1-32 ura4-D18 SPAC212.07*::*ura4*$^{+}$ Δ*mst2*::*hphMX6* | 7A, 7B | This study |
| **Bio7068** | $h^{+}$ *leu1-32 ura4-D18 SPAC212.07*::*ura4*$^{+}$ Δ*mst2*::*hphMX6* Δ*mrc1*::*kanMX6* | 7A, 7B | This study |
| **Bio7080** | $h^{?}$ *leu1-32 ura4-D18 SPAC212.07*::*ura4*$^{+}$ Δ*mst2*::*hphMX6 mrc1*Δ*782-879-3FLAG*::*kanMX6* (Δ*HBS*) | 7A, 7B | This study |
| **Bio7626** | $h^{-}$ *leu1-32 ura4-D18 ade6-m210 his2* Δ*K*::*ura4*$^{+}$Δ*mst2*::*natMX6* | 7C, 7D | This study |
| **Bio7733** | $h^{-}$ *leu1-32 ura4-D18 ade6-m210 his2* Δ*K*::*ura4*$^{+}$Δ*mst2*::*natMX6* Δ*mrc1*::*kanMX6* | 7C, 7D | This study |
| **Bio7734** | $h^{-}$ *leu1-32 ura4-D18 ade6-m210 his2* Δ*K*::*ura4*$^{+}$Δ*mst2*::*natMX6 mrc1*Δ*782-1019-5FLAG*::*kanMX6* (Δ*HBSC*) | 7C, 7D | This study |
| **AH109** | *MATa, trp1-901, leu2-3, 112, ura3-52, his3-200, gal4* Δ *, gal80* Δ *,LYS2::GAL1*$_{UAS}$*-GAL1*$_{TATA}$*-HIS3, MEL1, GAL2*$_{UAS}$*-GAL2*$_{TATA}$*-ADE2, URA3::MEL1*$_{UAS}$*-MEL1*$_{TATA}$*-lacZ* | 6B, 6C | The Matchmaker two-hybrid system 3 (Clontech) |
