## Supplementary material for "Mrc1^Claspin^ is essential for heterochromatin maintenance in *Schizosaccharomyces pombe*": Table_S2

**Table S2: Primer list**

| Name | Sequence (5' − 3') | Figure |
|---|---|---|
| ***ade6*****+** **fw** | GTAGTACGCAGTTTAGACGG | 1E, 1F, 3C, 3D |
| ***ade6*****+** **rv** | GAGCACGCTGTTGAATTGAG | |
| ***act1*****+** **fw** | TGGCTCTGGTATGTGCAAAG | 1E, 3C, 4B |
| ***act1*****+** **rv** | AGCTTCATCACCAACGTAGG | |
| ***tlh1*** **fw** | GTTGGCTAATTATGGACTCTC | 3C, 3D, 5A-5E, 7B |
| ***tlh1*** **rv** | CACAATTTGCATATGTTGCGC | |
| ***SPAC212.08*** **fw** | GGAGGACCTAGATTGGTTAAC | 3C, 3D, 5A-5E, 7B |
| ***SPAC212.08*** **rv** | CACAGCTAAAGTAATCAGCAC | |
| ***ura4*****+** **fw** | GAATGGTTTGAGAAGCATACC | 4D, 7B, 7D, S1B |
| ***ura4*****+** **rv** | GAGTACGATATTGCTGTCCC | |
| ***dg*** **fw** | CTGCGGTTCACCCTTAACATC | 4B, 5A-5E |
| ***dg*** **rv** | CAACTGCGGATGGAAAAAGT | |
