## Supplementary material for "Mrc1^Claspin^ is essential for heterochromatin maintenance in *Schizosaccharomyces pombe*": Table_S3

**Table S3: Plasmid list**

| Plasmid No. | Name | Description | Origin |
|---|---|---|---|
| **pKT880** | pGBKT7 | *Gal4-BD, TRP1, cm$^r$* | our stock |
| **pKT388** | pGADT7 | *Gal4-AD, LEU2, amp$^r$* | our stock |
| **pKT3** | pGBT9-mrc1 | *Gal4-BD-mrc1, TRP1, amp$^r$* | This study |
| **pKT67** | pGBKT7-mrc1-N | *Gal4-BD-mrc1(1-503), TRP1, cm$^r$* | This study |
| **pKT68** | pGBKT7-mrc1-M | *Gal4-BD-mrc1(248-756), TRP1, cm$^r$* | This study |
| **pKT69** | pGBKT7-mrc1-C | *Gal4-BD-mrc1(500-1019), TRP1, cm$^r$* | This study |
| **pKT3216** | pGADT7-clr1 | *Gal4-AD-clr1, LEU2, amp$^r$* | This study |
| **pKT3217** | pGADT7-clr2 | *Gal4-AD-clr2, LEU2, amp$^r$* | This study |
| **pKT3218** | pGADT7-clr3 | *Gal4-AD-clr3, LEU2, amp$^r$* | This study |
| **pKT3219** | pGADT7-mit1 | *Gal4-AD-mit1, LEU2, amp$^r$* | This study |
